# Nitrates increase abscisic acid levels to regulate haustoria formation in the parasitic plant Phtheirospermum japonicum

**DOI:** 10.1101/2021.06.15.448499

**Authors:** Anna Kokla, Martina Leso, Xiang Zhang, Jan Simura, Songkui Cui, Karin Ljung, Satoko Yoshida, Charles W. Melnyk

## Abstract

Parasitic plants are globally prevalent pathogens that withdraw nutrients from their host plants using an organ known as the haustorium. Some, the obligate parasites are entirely dependent on their hosts for survival, whereas others, the facultative parasites, are independent of their hosts and infect depending on environmental conditions and the presence of the host. How parasitic plants regulate their haustoria in response to their environment is largely unknown. Using the facultative root parasite *Phtheirospermum japonicum*, we found that external nutrient levels modified haustorial numbers. This effect was independent of phosphate and potassium but nitrates were sufficient and necessary to block haustoria formation. Elevated nitrate levels prevented the activation of hundreds of genes associated with haustoria formation, downregulated genes associated with xylem development and increased levels of abscisic acid (ABA). Enhancing ABA levels independently of nitrates blocked haustoria formation whereas reducing ABA biosynthesis allowed haustoria to form in the presence of nitrates suggesting that nitrates mediated haustorial regulation in part via ABA production. Nitrates also inhibited haustoria formation and reduced infectivity of the obligate root parasite *Striga hermonthica*, suggesting a more widely conserved mechanism by which parasitic plants adapt their extent of parasitism according to nitrogen availability in the external environment.

## Introduction

Parasitic plants make up ~1% of all angiosperm species; some of which are devastating agricultural weeds that cause major agricultural loses each year (De Groote et al., 2007; Heide-Jørgensen, 2008; Rodenburg et al., 2016). Parasitic plants can range from obligate parasites that completely depend on their host for survival to facultative parasites that can survive without a host but parasitize when conditions are suitable (Heide-Jørgensen, 2008; Spallek et al., 2013). Despite differences in their lifestyle, all parasitic plants form an invasive organ termed the haustorium (Kuijt, 1969) through which they penetrate the host and uptake water, nutrients, RNA and hormones (Barkman et al., 2007; Kuijt, 1969; Kokla and Melnyk, 2018; Spallek et al., 2017; Shahid et al., 2018).

Many parasitic plants, particularly the obligate parasites, require perception of host-exuded compounds such as strigolactones to initiate germination. Perception of a second host-derived compound, known as haustorium inducing factors (HIFs), initiates haustorium formation in both obligate and facultative parasites. The first identified HIF was 2,6-dimethoxy-1,4-benzoquinone (DMBQ), originally isolated from root extracts of infected sorghum plants. DMBQ can induce haustoria formation even in the absence of a host (Chang and Lynn, 1986) in a wide range of parasitic plants. In the facultative parasitic plant *Phtheirospermum japonicum*, perception of a nearby host via HIFs is followed by cell expansion and division at the haustorium initiation site, forming the characteristic swelling of the pre-haustorium. Later, the developing haustorium attaches to the host and starts penetrating to reach the vascular cylinder of the host. Once the haustorium has reached the host’s vasculature, it starts forming a xylem connection between itself and the host known as the xylem bridge (Cui et al., 2016; Ishida et al., 2016; Wakatake et al., 2018; Heide-Jorgensen and Kuijt, 1995).

Despite recent advances in our understanding of haustorium development, little is known about how environmental conditions affect plant parasitism. Nutrient availability is an important factor affecting plant parasitism, for example, infestations of the agriculturally devastating obligate parasite *Striga* are often associated with poor soil fertility (Mwangangi et al., 2021). Low soil fertility is thought to impede host defences and exacerbate the damaging effects of infection (Mwangangi et al., 2021). In addition, low nutrient levels in the soil, particularly phosphate, promotes host secretion of strigolactones which enhances *Striga* germination and infections levels. Improving soil fertility can reduce the production of germination stimulants while also improving host defences and host tolerance (Jamil et al., 2012; Yoneyama et al., 2007a, 2007b; Sun et al., 2014; Mwangangi et al., 2021). However, nutrients might also have effects on the parasite beyond germination. For instance, application of certain nitrate compounds reduced *Striga* shoot development (Igbinnosa et al., 1996) whereas *Phtheirospermum japonicum* required nutrient starvation to efficiently infect its hosts *in vitro* (Cui et al., 2016; Ishida et al., 2016; Spallek et al., 2017) and *Rhinanthus minor* growth was inhibited in the presence of high phosphorus (Davies and Graves, 2000). Together, these data suggest that nutrients might play a role beyond improving host fitness or reducing parasite germination.

Nutrient availability affects many aspects of plant development including germination, root growth, shoot growth and flowering (Zhang and Forde, 2000; Alboresi et al., 2005; Castro Marín et al., 2011). High nitrate levels generally promote shoot growth and repress root growth, in part, through the action of plant hormones. In *Arabidopsis thaliana*, rice, maize and barley, nitrates increase cytokinin levels which move to the shoot meristems to promote cell divisions and growth (Samuelson and Larsson, 1994; Takei et al., 2001, 2004; Kamada-Nobusada et al., 2013; Landrein et al., 2018). Nitrates also inhibit auxin transport and modifies auxin response to promote root initiation but inhibit root elongation (Vidal et al., 2010). ABA too plays a role; nitrate treatments increase ABA levels in *Arabidopsis* root tips (Ondzighi-Assoume et al., 2016) whereas ABA signaling is required for the inhibitory effects of high nitrates on root growth (Signora et al., 2001). However, the mechanisms through which nutrient availability affects plant parasitism remains unknown.

Here, we show that nutrient rich soils greatly reduce both root size and haustorial density in *Phtheirospermum japonicum*, and this effect is dependent specifically on nitrate concentrations. Nitrate application blocked gene expression changes associated with haustoria formation and modified xylem patterning in the root. Nitrates increased ABA levels and activated ABA responsive genes expression. Treating with ABA reduced haustoria initiation whereas inhibiting ABA biosynthesis could reduce the inhibitory effects of nitrates. Finally, we investigated the effects of nutrients in *Striga hermonthica* and found that similar to *Phtheirospermum japonicum*, nutrients decreased haustoria formation rates and infection rates, and this effect was specific to nitrates.

## Results

### Nitrate inhibits haustoria development

Low nutrients are important for efficient *Striga* infestations and successful *Phtheirospermum japonicum in vitro* infections (Oswald, 2005; Ishida et al., 2011; Cui et al., 2016; Spallek et al., 2017). We tested whether successful *Phtheirospermum-Arabidopsis* soil infections also required low nutrients by treating nutrient poor 50:50 soil:sand with or without fertilizer. *Phtheirospermum* shoot weights and heights were similar in both treatments, but root masses and haustorial density were higher in nutrient poor conditions (Fig.1A-C; Fig.S1A-C). To better understand the basis for reduced haustoria in nutrient rich conditions, we grew 4-5-day old *Phtheirospermum* seedlings *in vitro* on water-agar or half-strength Murashige and Skoog medium (½MS)-agar (Fig.S1E). Similar to fertilized soil, *Phtheirospermum-Arabidopsis* infections on ½MS-agar formed substantially fewer infections than those on water-agar (Fig.1D). On ½MS, pre-haustoria were initiated but they did not penetrate the host or form xylem bridges (Fig.1E, I). To identify the compound(s) that caused haustoria arrest, we tested three of the major macroelements found in MS plus one macrolement found in Gamborg’s B5 medium at similar concentrations as those found in ½MS or Gamborg’s medium. Agar media containing phosphate (KH_2_PO_4_ or NaH_2_PO_4_) or potassium (KH_2_PO_4_) had no effect on haustoria formation, but agar media containing nitrate (NH_4_NO_3_ or KNO_3_) inhibited *Phtheirospermum-Arabidopsis* infections and xylem bridge formation similar to ½MS (Fig.1D, E, I). Infections on ½MS lacking nitrates did not affect haustoria or xylem bridge formation (Fig.1D, E, I; Fig.S1D, F, G) indicating that nitrates were sufficient and necessary to block infections. Nitrate application led to a reduction of haustoria and xylem bridge formation in a wide range of concentrations from 50 μM to 20.6 mM (Fig.1F, G; Fig.S1D, F, G). To test whether nitrate blocked infection by inhibiting the parasite or strengthening the host, we applied NH_4_NO_3_ or ½MS to *Phtheirospermum* growing alone in the presence of the haustoria inducting factor DMBQ. Combining DMBQ with water or ½MS lacking nitrate formed similar numbers of pre-haustoria, whereas combining DMBQ with NH_4_NO_3_ or ½MS greatly reduced pre-haustoria formation (Fig.1H, I) suggesting the effect of nitrate on haustoria initiation was specific to the parasite.

**Fig.1.**
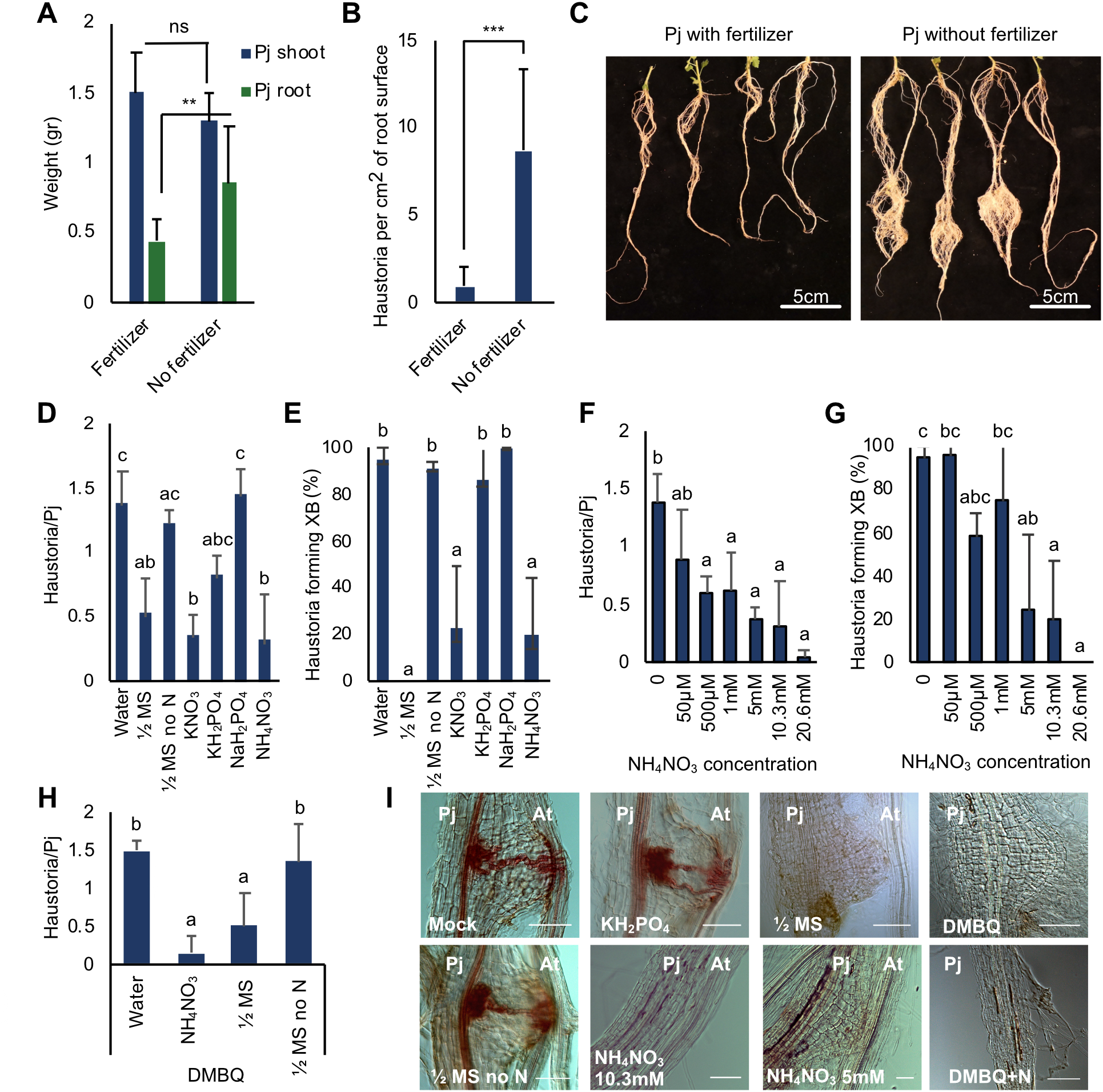
Nitrogen inhibits *Phtheirospermum* haustoria formation. (A-C) *Phtheirospermum* shoot and root weight during *Arabidopsis* infection with and without fertilizer application.(B) Haustoria numbers per cm^2^ *Phtheirospermum* root surface during *Arabidopsis* infection with and without application of fertilizer. (D-E) Average number of haustoria per *Phtheirospermum* seedling and xylem bridge formation percentage in *in vitro* infection assays with *Arabidopsis* (Col-0) as the host with various nutrient treatments (½MS, ½MS no N, 20.6 mM KNO_3_, 10.3 mM NH_4_NO_3_, 0.62 mM KH_2_PO_4_ or 1.9 mM NaH_2_PO_4_) and (F-G) with a range of NH_4_NO_3_ concentrations (20.6 mM to 50 μM). (H) Average number of haustoria *per Phtheirospermum* seedling with 10μM DMBQ and nutrient application (½MS, ½MS no N or 10.3 mM NH_4_NO_3_). (I) Brightfield images of *Phtheirospermum* haustoria during *Arabidopsis in vitro* infections with nutrient treatments. (A-I) Bars represent mean ± SD (ANOVA P<0.05). **P<0.001, ***P<0.0001, Student’s t-test, two tailed. Scale bars 50 μm for (I).

### Haustoria formation induces widespread transcriptional changes

To further investigate how nitrate availability affected haustoria formation in *Phtheirospermum*, we performed a time course RNAseq experiment of *Phtheirospermum* infecting *Arabidopsis* Col-0 in an *in vitro* infection assay with water-agar or NH_4_NO_3_ treatments. Nitrate treatments increase cytokinin levels in *Arabidopsis*, rice, maize and barley (Samuelson and Larsson, 1994; Takei et al., 2001, 2004; Kamada-Nobusada et al., 2013) so we also included infections with 6-benzylaminopurine (BA), a synthetic cytokinin, to test similarities between NH_4_NO_3_ and BA transcriptional responses. *Phtheirospermum* and *Arabidopsis* were aligned at time 0 to help synchronize infections (Fig.S1E) and tissues surrounding the root tips where haustoria normally emerge were collected at 0,12, 24, 48, 72 hours post infection (hpi) for the water-only treatment and 0,12, 24 hpi for the NH_4_NO_3_ and BA treatments (Fig.2A). With water-only treatments, we observed an increasing number of differentially expressed genes as the infection progressed (Fig.2C). Co-expression analyses enabled us to classify genes into 8 clusters with distinct expression patterns during haustorium formation (Fig.2B, D, TableS1). Cluster 2, 3 and 8 whose gene expression peaked at early stages of haustoria formation (12 and 24hpi) had an over representation of genes that belong to Gene Ontology enrichment (GO) categories related to transcription, translation, signaling processes and cell expansion/replication (Fig.S2A). Cluster 4, 5 and 7 whose gene expression peaked at later time points in haustorium formation (48 and 72hpi) had an over representation of genes that belong to GO categories related to response to oxidative stress, cytokinin metabolic process, fatty acid biosynthetic process, lignin, sucrose and carbohydrate metabolism (Fig.S2A). We looked at the expression of individual genes in our transcriptome and identified an upregulation of auxin-related *PjYUC3, PjLAX1, PjPIN9*, and cambium-related *PjWOX4*, genes whose expression has been previously observed to increase during *Phtheirospermum* infections (Ishida et al., 2016; Wakatake et al., 2018, 2020) (Fig.2E). Genes associated with cytokinin metabolism (*PjCKX3, PjCKX1,PjLOG8*) and cell wall remodeling (*PjPMEI*9) were upregulated as well (Fig.2E).

**Fig. 2.**
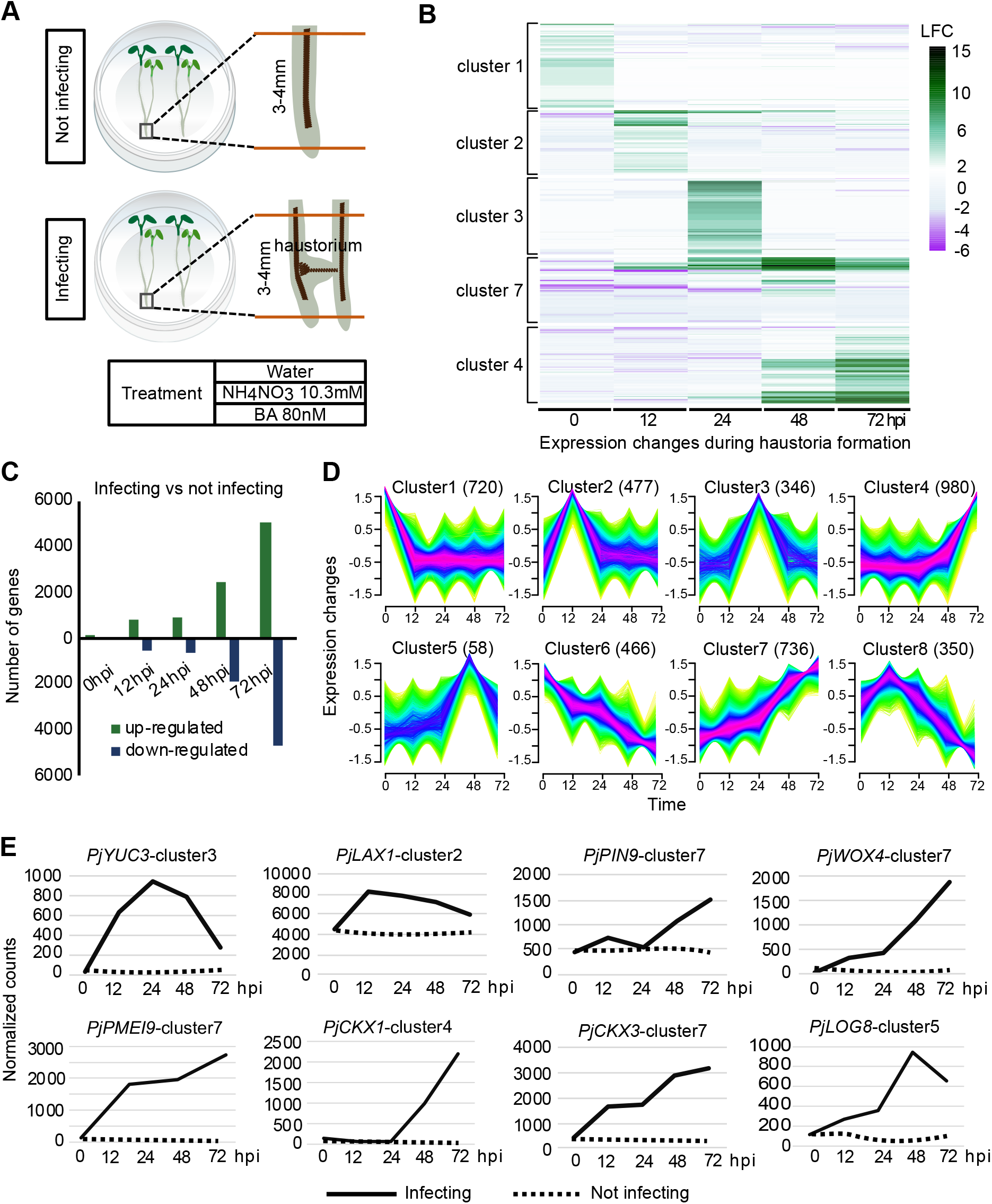
Transcriptomic changes during haustorium formation. (A) Illustration describing the experimental set-up for the RNAseq. (B) Heatmap of the log2 fold change of gene subsets that belong to five co-expression clusters over five time points in the water-only RNAseq treatment between *Phtheirospermum* infecting and not infecting. (C) Number of genes differentially expressed over five time points in the water-only RNAseq treatment comparing *Phtheirospermum* infecting versus not infecting. (D) Clustering of DE genes in the water-only RNAseq treatment of *Phtheirospermum* infecting *Arabidopsis* over five time points based on their co-expression patterns; the number next to the dash represents the number of genes in the respective cluster. (E) Normalized counts of *PjYUC3, PjLAX1, PjPIN9, PjWOX4, PjPMEI9, PjCKX1, PjCKX3, PjLOG8* over 5 time points in the water-only RNAseq treatment for *Phtheirospermum* infecting and not infecting.

### Nitrate inhibits genes associated with early haustorial development

Nitrate prevented haustoria formation (Fig.1) so we looked at when this block occurs transcriptionally. We compared infected NH_4_NO_3_ samples with not infected NH_4_NO_3_ samples and found fewer than 70 differentially expressed genes at any time point (Fig.S3A). Less than 20 of these genes at each time point were also differentially expressed during water-only infections. This low number suggested that nitrates blocked the vast majority of haustorial-related gene activation and stopped haustorial formation early during infection. We next compared the transcriptional differences between water-only infections and NH_4_NO_3_ infections and found between 4000 and 6000 genes were expressed differently between treatments at each time point (Fig.S3B). Genes that were upregulated during successful haustoria formation were not activated in the NH_4_NO_3_ treatment including *PjYUC3, PjWOX4* and *PjPMEI9* (Fig.3A, B), consistent with nitrate acting early to block haustoria induction. We looked at the not infected NH_4_NO_3_ datasets to see which genes might be responsible for an early block in haustorial-related genes. We found a GO enrichment for cell wall and lignin-related genes downregulated in the not infecting NH_4_NO_3_ treatment compared to the not infecting water treatment (Fig.S4A). Genes downregulated included xylem-related *PjXCP1, PjLAC11, PjIRX3, PjCESA4, PjPRX66* and quinone perception related genes *PjCADL2* and *PjCADL4* (Laohavisit et al., 2020) (Fig.3B, Fig.S3G,H). Cytokinin-related GOs were also enriched in the genes downregulated by nitrate (Fig.S4A) and we found no substantial overlap between differentially expressed genes in the BA-treated and NH_4_NO_3_-treated not infected samples (Fig.S5I). Together, these data suggested that nitrate did not induce cytokinin response and that nitrate blocked the infection process at an early stage.

**Fig. 3.**
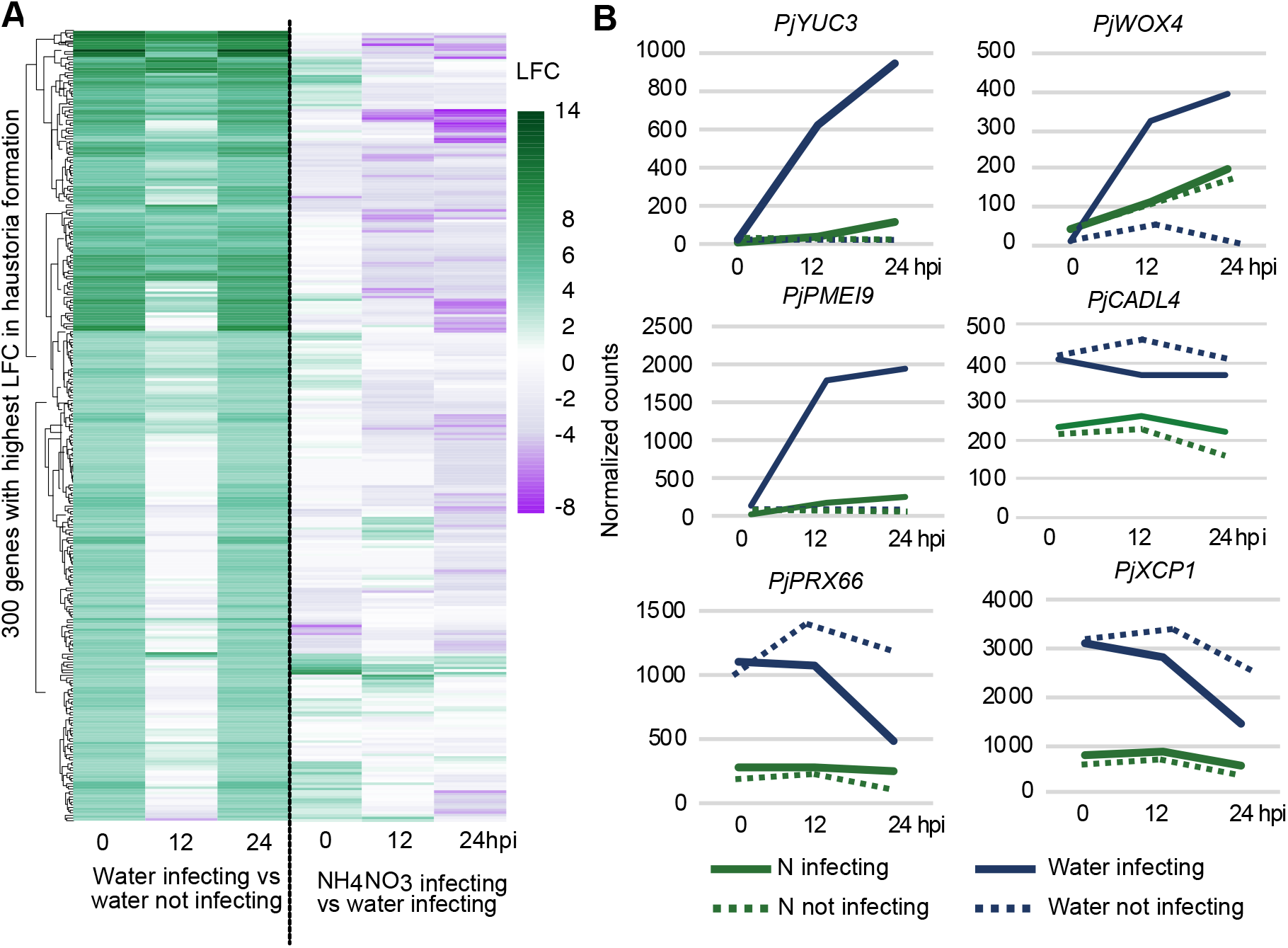
NH_4_NO_3_ treatment modifies *Phtheirospermum* gene expression. (A) Heatmap of 300 genes with the highest log2 fold change during haustoria formation shown over three time points in the water infecting versus water not infecting and NH_4_NO_3_ infecting vs water infecting RNAseq treatments in *Phtheirospermum.* (B) Normalized counts of *PjYUC3, PjWOX4, PjPMEI9, PjXCP1, PjCADL4, PjPRX66* over three time points shown for *Phtheirospermum* infecting and not infecting in the NH_4_NO_3_ and water treatment.

### Nitrate increases ABA levels in *Phtheirospermum*

To further investigate how nitrate leads to the arrest of haustoria formation, we performed hormonal profiling on 50-day-old *Phtheirospermum* roots that have been infecting *Arabidopsis* for 10 days with and without nitrate treatment. Cytokinin levels increased in successful infections, similar to a previous study (Spallek et al., 2017), but we observed no other hormones substantially induced only by infection (Fig.4A, B). However, ABA and salicylic acid (SA) were significantly increased in *Phtheirospermum* roots treated with nitrate in both not infected and infected samples compared to water alone (Fig.4A). We further quantified ABA levels in 20-day-old *Phtheirospermum* whole seedlings that have been infecting *Arabidopsis* for 10 days. Similar to older root tissues, ABA levels increased in both infecting and not infecting nitrate treated *Phtheirospermum* seedlings compared to water-only treatments (Fig.4C). ABA levels in the host *Arabidopsis* Col-0 were not significantly affected by nitrate treatments but instead were increased during *Phtheirospermum* infection (Fig.4D). This increase was dependent on host ABA biosynthesis since the increase was blocked in the ABA biosynthesis mutant *aba2-1* (Fig.4D). Cytokinin moves from *Phtheirospermum* to *Arabidopsis* during infections (Spallek et al., 2017) but we found no evidence that ABA moved from parasite to host since *Arabidopsis* ABA levels in *aba2-1* infections were similar to not infected *aba2-1* plants (Fig.4C-D). Consistent with our hormone quantification experiments, the transcriptome analysis revealed that *Phtheirospermum* genes homologous to *Arabidopsis* ABA responsive genes had increased expression levels in the NH_4_NO_3_ treatment compared to the water treatment for both infected and not infected tissues (Fig.4E, Fig.S4B). This expression pattern was not seen when comparing the same genes to the BA treatment or non-nitrate treated samples (Fig.S4C, S5A, D). Most cytokinin related genes were not differentially expressed in the NH_4_NO_3_ treatment compared to the water treatment - with some exceptions – further supporting our finding that NH_4_NO_3_ treatment in *Phtheirospermum* does not induce widespread cytokinin response (Fig.4F, Fig.S4D). However, cytokinin related genes were differentially expressed during later time points in water treatment infections implicating cytokinin in successful haustorium development (Fig.S5E). To test whether the *Phtheirospermum* genes homologous to *Arabidopsis* ABA responsive genes were ABA responsive, we selected four genes and found by qPCR that their transcript levels were increased by exogenous ABA (Fig.S5B). We tested the expression levels of the same genes in *Phtheirospermum* grown on various soil:sand ratios and found that the expression levels of these genes were higher in the 100% soil (high nutrient availability) than 25%soil (low nutrient availability)(Fig.S5C), suggesting that soil grown plants show similar ABA responses to high nitrates. SA levels also increased during nitrate treatment (Fig.4A) but most *Phtheirospermum* genes homologous to *Arabidopsis* SA-related genes were not differentially expressed in the NH_4_NO_3_ treatment compared to the water treatment (Fig.S5F). Together, these data suggest that nitrates specifically induced ABA levels and increased ABA response.

**Fig. 4.**
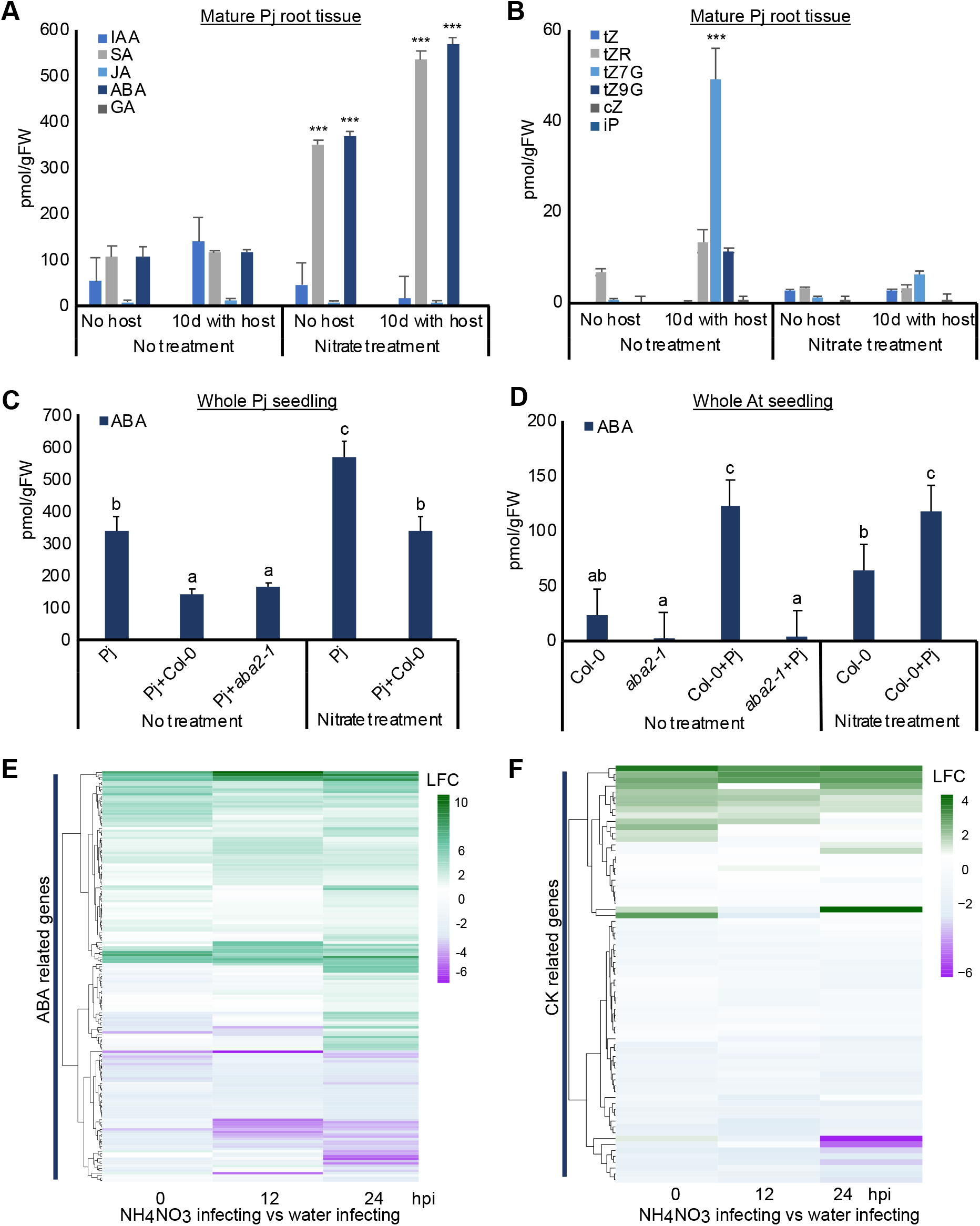
ABA levels increase during nitrate treatment. (A-B) Hormonal quantification of *Phtheirospermum* roots treated with 10.3 mM NH_4_NO_3_. (C-D) Hormonal quantification of *Phtheirospermum* whole seedlings and *Arabidopsis* (Col-0, *aba2-1*) whole seedlings treated with 10.3 mM NH_4_NO_3_. (E) Heatmap of the log2 fold change of 170 genes homologous to *Arabidopsis* ABA responsive genes shown over three time points in the NH_4_NO_3_ infecting vs the water infecting RNAseq treatment in *Phtheirospermum*. (F) Heatmap of the log2 fold change of 72 genes homologous to *Arabidopsis* cytokinin related (signaling-biosynthesis-metabolism) genes shown over three time points in the NH_4_NO_3_ infecting vs the water infecting RNAseq treatment in *Phtheirospermum*. (A-D) Bars represent mean ± SD (ANOVA P<0.05). ***P<0.0001, Student’s t-test, two tailed.

### ABA affects haustoria formation

To investigate the role of ABA on haustorium formation in *Phtheirospermum*, we applied ABA exogenously using *in vitro* infection assays. ABA treated plants formed less haustoria than water-only treated plants with some haustoria not forming xylem bridges (Fig.5A, B, I). The application of fluridone, a chemical that inhibits ABA biosynthesis, significantly reduced xylem bridge formation but had no effect on haustorial numbers (Fig.5A, B, E, F, I). We reasoned that if nitrates induce ABA to repress haustoria, we could overcome the inhibitory effects of nitrate by blocking ABA biosynthesis with fluridone. Indeed, combining ½MS treatments with fluridone increased haustoria numbers compared to the ½MS treatment but they remained intermediate to the water-only treatment (Fig.5E, F, I). *Phtheirospermum* infecting *aba2-1* or *aba1-1C* did not have differences in haustoria and xylem bridge formation, suggesting that altering host ABA biosynthesis or signaling did not affect parasitism (Fig.S6A, B). SA levels were also induced by nitrates in *Phtheirospermum* (Fig.4) and we observed that exogenous application of SA decreased haustorial numbers but did not affect xylem bridge formation (Fig.5C, D, I).

**Fig.5.**
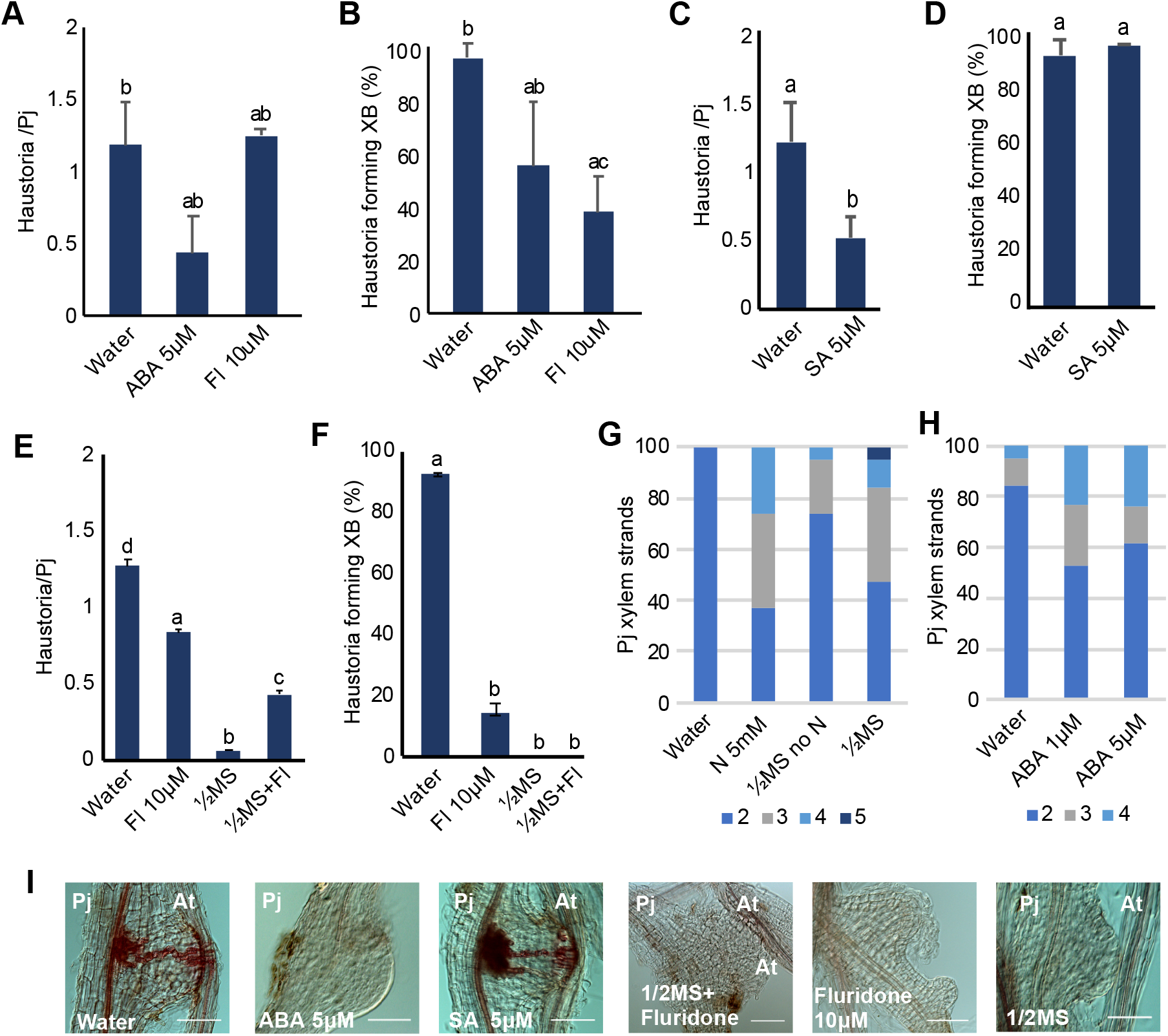
ABA represses *Phtheirospermum* haustoria formation. (A-F) Average number of haustoria per *Phtheirospermum* seedling and xylem bridge formation percentage in *in vitro* infection assays with *Arabidopsis* (Col-0) as the host treated with 5 μM ABA, 10 μM fluridone (Fl), 5 μM SA,1/2MS or 1/2MS no N. (G-H) Number of lignified xylem strands at 2 mm from the root tip in *Phtheirospermum* seedlings treated with 5 mM NH_4_NO_3_, 1/2MS, 1/2MS no N,1μM ABA or 5μM ABA. (I) Brightfield images of *Phtheirospermum* haustoria during *Arabidopsis in vitro* infection under 5μM ABA, 10 μM fluridone, 1/2MS, 1/2MS + 10 μM fluridone or 5μM SA treatments. (A-F) Bars represent mean ± SD (ANOVA P<0.05). Scale bars 50 μm for (I).

Since ABA is important for various developmental processes including root xylem formation (Ramachandran et al., 2021), we analyzed the *Phtheirospermum* transcriptome under low nitrate conditions and found that some *Phtheirospermum* genes homologous to *Arabidopsis* ABA responsive genes had increased expression levels during successful infection (Fig.S5D). Exogenous ABA treatments did not increase xylem bridge numbers or size (Fig.5, S6) but treatment with fluridone blocked xylem bridge formation, consistent with ABA playing a role during xylem bridge formation. Exogenous ABA application to *Phtheirospermum* was previously shown to enhance the number of differentiating xylem strands in primary root tips (Ramachandran et al., 2021). We repeated this assay but also included nitrate treatments on *Phtheirospermum* seedlings. ABA, NH_4_NO_3_ and ½MS all had a similar phenotype of increased xylem strand differentiation (Fig.5G, H). These data suggested that ABA played additional roles in both xylem bridge formation and also in modulating xylem patterning in response to nitrate levels.

### Nitrates affect *Striga* infection rates

Previous field studies showed that *Striga* infection is decreased after nitrate application (Jamil et al., 2012; Cechin and M. C. Press, 1993). We investigated the effect of nutrients using *in vitro Striga hermonthica* infection assays. In the presence of NH_4_NO_3_, KNO_3_, KH_2_PO_4_ or NaH_2_PO_4_ *Striga* infection rates were not significantly decreased at two weeks after infection (Fig.6A). However, at four weeks after infection nitrate application lead to a significant decrease in *Striga* infection rates (Fig.6A, D). *Striga* development was also hindered in the presence of nitrates where the appearance of plants with 3-5 leaf pairs and more that 6 leaf pairs were decreased compared to the water treatment (Fig.6B). We tested whether this effect was mediated by improved host fitness or reduced *Striga* infectivity by treating *Striga* with DMBQ in the presence of nitrate. Prehaustoria formation by DMBQ was significantly reduced in the presence of NH_4_NO_3_ (Fig.6C) suggesting that like *Phtheirospermum*, early *Striga* haustoria formation is inhibited by high nitrates. *Striga* is highly resistant to ABA (Fujioka et al., 2019), but nonetheless we tested exogenous application of ABA or fluridone and found they did not have an effect on *Striga* haustoria formation (Fig.S7).

**Fig.6.**
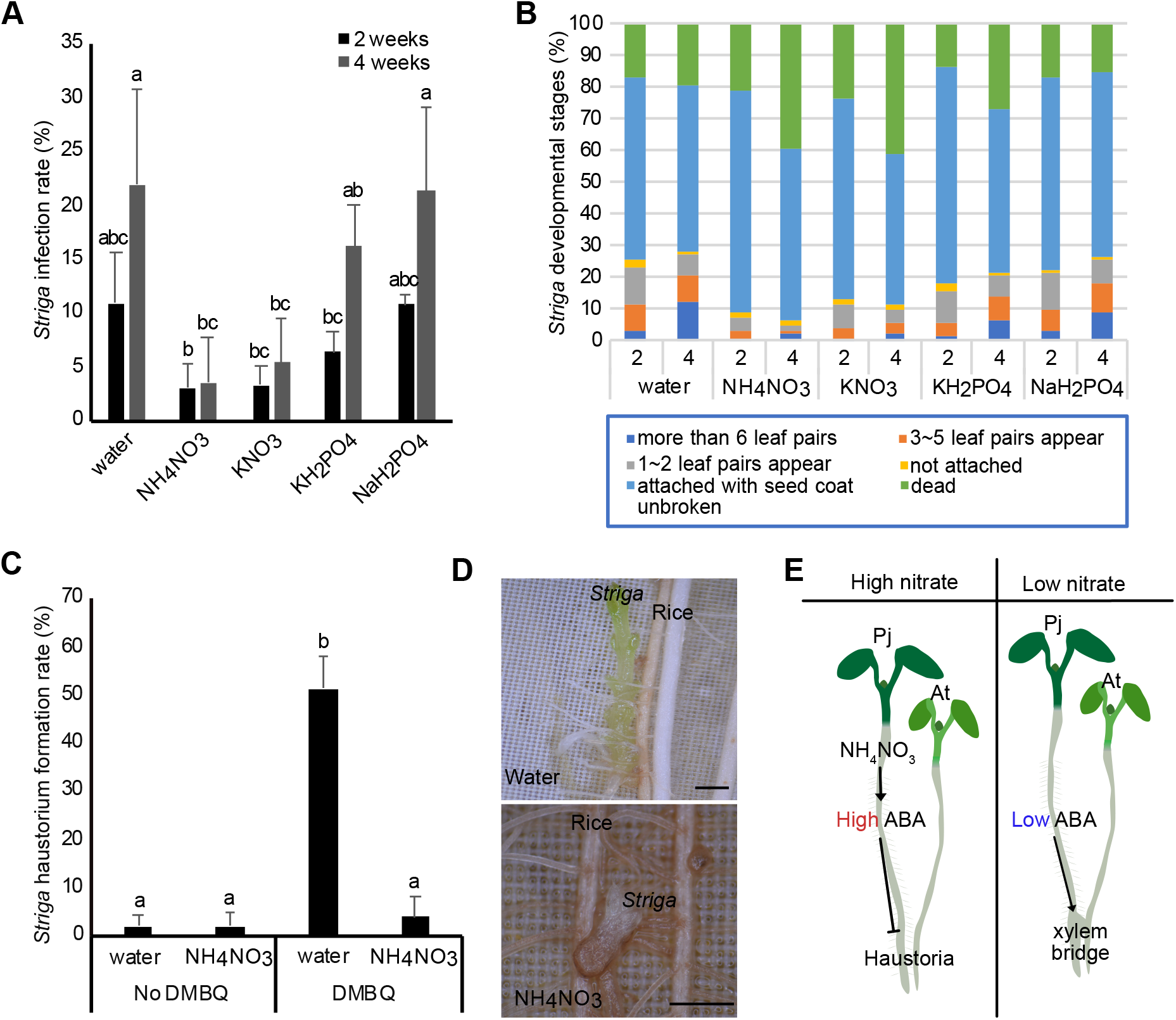
NH_4_NO_3_ inhibits *Striga* infection rates. (A) *Striga* infection rate at two and four weeks after infection with rice as a host under nutrient treatments (20.6 mM KNO_3_, 10.3 mM NH_4_NO_3_, 0.62 mM KH_2_PO_4_, 1.9 mM NaH_2_PO_4_). (B) Effect of nutrient treatments on *Striga* development at two and four weeks after infection. (C) Effect of NH_4_NO_3_ treatment on *Striga* haustorium formation induced by 1 μM DMBQ. (D) Brightfield images of *Striga* infecting rice at 2 weeks after infection. (E) Graphical representation of a putative model of nitrate-ABA mediated haustoria regulation. (A,C) Bars represent mean ± SD (ANOVA P<0.05). Scale bars 1 mm for (D).

## Discussion

### The effect of nitrogen on parasitism

Here, we describe a mechanism whereby external nitrate levels regulate haustoria formation in the facultative root parasite *Phtheirospermum japonicum* (Fig.6E). This effect did not occur with phosphate or potassium and instead appeared highly specific to nitrate in micromolar concentrations (Fig.1F, G). Increased nitrogen supply to *Medicago sativa* also reduced *Phtheirospermum japonicum* parasitism (Irving et al., 2019), consistent with our results, and we propose that local nitrogen supply at the site of infection has a suppressive effect upon the parasite rather than the host. *Striga* haustoria formation and infection rates were also inhibited by nitrate (Fig.6A-C) which is consistent with previous observations that external nitrates reduce *Striga* growth (Cechin and M. C. Press, 1993; Igbinnosa et al., 1996). However, our results point to an earlier role for nitrates by preventing haustoria to develop beyond the initial swell in the presence of the HIF DMBQ (Fig.1H, Fig. 6C). Nitrate might block HIF perception in the root, or alternatively, starvation induces competency for HIF perception and haustoria elongation. As such, the observed reduction in *Striga* infestations in nutrient rich fields (Jamil et al., 2012) could be from a combination of reduced germination stimulants, reduced haustoria formation and a suppression of *Striga* growth (Cechin and M. C. Press, 1993; Igbinnosa et al., 1996). Our results also suggest a conserved role for nitrates regulating haustoria in both facultative and obligate Orobanchaceae family members.

Beyond parasitism, nitrate has strong effects upon plant root architecture and organogenesis. In *Arabidopsis*, mild nitrogen deficiency enhances lateral root elongation, whereas uniform high nitrate levels repress lateral root development (Bouguyon et al., 2015; Araya et al., 2016; Zhang et al., 1999). In nodulating plants, high nitrogen levels in the environment repress nodule formation in *Medicago truncatula*, soybean and alfalfa through a regulatory mechanism involving multiple hormones and peptides (Schultze and Kondorosi, 1998; Carroll et al., 1985; Caba et al., 1998; Reid et al., 2011; Gautrat et al., 2019). Our data and these results suggest a common regulatory theme whereby low nitrate levels promote organ growth to uptake additional nutrients, whereas high nitrate levels repress organ growth to avoid unnecessary resources spent on nutrient acquisition. These findings might imply that *Phtheirospermum* and *Striga* are specifically looking to acquire nitrate from their hosts, however, it is likely that multiple nutrients are obtained but nitrate is used as an environmental regulatory cue.

### The role of hormones in nitrogen induced haustoria regulation

In species like *Arabidopsis*, maize, rice and barley, high nitrates increase cytokinin levels that are important for root development and shoot growth (Samuelson and Larsson, 1994; Kamada-Nobusada et al., 2013; Takei et al., 2004, 2001). Cytokinin levels were strongly induced by *Phtheirospermum* infection (Fig.4), consistent with previous findings (Spallek et al., 2017), but we found no evidence that nitrate itself induced cytokinin levels or induced cytokinin response in *Phtheirospermum*. Notably, nitrate treatment of *Lotus japonicus* inhibited cytokinin biosynthesis, reduced cytokinin levels and reduced nodule formation (Lin et al., 2020). In *Phtheirospermum*, nitrate treatment also reduced cytokinin levels and cytokinin response compared to water treatment (Fig.4B, Fig.S4). Thus, both *Phtheirospermum* and *Lotus* appear to use nitrates to suppress haustoria and nodules independently of cytokinin biosynthesis, whereas cytokinin production instead indicates successful symbiosis. This situation differs from most other flowering plants and might be a convergent strategy to use cytokinin to signal successful organogenesis rather than nutrient abundance.

We observed that nitrate increased ABA levels in *Phtheirospermum* independently of infection. ABA levels are known to increase in both *Rhinanthus minor* and *Cuscuta japonica*, as well as their hosts, after infection (Jiang et al., 2004; Furuhashi et al., 2014). *Striga* parasitism commonly induces symptoms in the host mimicking drought stress and increases host ABA levels in tomato and maize (Taylor et al., 1996; Frost et al., 1997). We too observed ABA levels increased in the host *Arabidopsis* upon parasitic plant infection, likely due to a stress or defense response rather than movement from parasite to host (Liao et al., 2016; Cheng et al., 2016; Van Gijsegem et al., 2017). Our treatments with exogenous ABA reduced haustoria numbers whereas treatment with the ABA biosynthesis inhibitor fluridone partially rescued the inhibition by nitrates, suggesting that ABA in part regulates haustoria (Fig.5). Other factors including SA or proteins known to affect lateral root or nodule formation likely also play a role. *Striga* and *Cuscuta* are highly insensitive to ABA (Fujioka et al., 2019; Li et al., 2015) and *Striga* did not respond to ABA in our assays (Fig.S7), indicating that these species and *Phtheirospermum* likely use additional mechanisms for nitrate-induced haustoria repression.

Our assays with nitrate and ABA revealed a dual role for ABA in *Phtheirospermum*. Low levels of ABA appeared important for xylem bridge formation, however, high levels from exogenous ABA treatment or nitrate application blocked haustoria formation (Fig.5). ABA treatment also produced haustoria that were underdeveloped or did not attach well, likely explaining the partial reduction in xylem bridge formation from ABA treatment (Fig.5). The situation in *Phtheirospermum* primary root tips differed since nitrate and ABA treatments induced early xylem differentiation (Fig.5) yet haustoria and surrounding tissues had reduced expression of xylem-related genes (Fig.3). These apparent differences between phenotype and expression might relate to differences in how the primary root and haustorium respond to ABA.

In nodulating plants, such as *Lotus japonicus, Trifolium repense* and *Medicago truncatula*, ABA acts as a negative regulator of nodules by repressing nod factor signaling and cytokinin responses (Ding et al., 2008; Suzuki et al., 2004; Tominaga et al., 2010). Exogenous application of ABA blocked the early stages of infection in *Lotus japonicus* (Suzuki et al., 2004), similar to the situation in *Phtheirospermum*. We propose that at least some parasitic plants and legumes share another common regulatory theme whereby ABA inhibits symbiotic organ formation, however, more work will be required to investigate these parallels including whether nitrates induce ABA in legumes and whether ABA inhibits HIF signaling in parasitic plants. In support of this idea, nitrate treatments reduced the expression of some of the *Phtheirospermum* quinone receptors that respond to DMBQ (Fig.3B, Fig.S3). Given that legumes and most parasitic plants are distantly related, it begs the question of whether such similar ABA and cytokinin regulatory features might be an important adaptation for symbiotic nutrient acquisition.

## Methods

### Plant materials and growth conditions

*Phtheirospermum* (Thunb.) Kanitz ecotype Okayama seeds were handled as described previously (Yoshida and Shirasu, 2009). *Arabidopsis* ecotype Columbia (Col-0) accession was used as *Arabidopsis* wild-type (WT). *Arabidopsis aba2-1* and *abi1-1C* were published previously (González-Guzmán et al., 2002; Umezawa et al., 2009). For *in vitro* germination, seeds were surface sterilized with 70% (v/v) EtOH for 20 minutes followed by 95%(v/v) EtOH for 5 minutes. The seeds were then sown on petri dishes containing ½MS medium (0.8% (w/v) plant agar, 1% (w/v) sucrose, pH 5.8). After overnight stratification in the dark and 4°C, the plants were transferred to 25°C long day conditions (16-h light:8-h dark and light levels 100 μmol m^-2^ s^-1^).

*Striga hermonthica* (Del.) Benth seeds were kind gifts provided by Dr A. G. T. Babiker (Environment and Natural Resources and Desertification Research Institute, Sudan). Rice seeds (*Oryza sativa L. subspecies japonica*, cvs Koshihikari) used in this study were originally obtained from National Institute of Biological Sciences (Tsukuba, Japan) and propagated in the Yoshida laboratory. The *Striga hermonthica* seeds were sterilized with a 20% (v/v) commercial bleach solution for 5 min and washed thoroughly with sterilized water on a clean bench. After that, these surface-sterilized *Striga* seed were placed in 9 cm petri dishes with moisturized glass fiber filter paper (Whatman GF/A) and preconditioned at 25 °C in the dark for 7 days. The preconditioned *Striga* seeds were treated with 10 nM Strigol (Hirayama and Mori, 1999) for 2 hours prior to rice-infection treatments. For haustorium induction assays, the preconditioned *Striga* seeds were treated with 10 nM Strigol at 25 °C for 1 day in the dark before starting incubation in various nutrient media with or without DMBQ and hormones for 24 hours in dark condition.

Rice seeds were de-husked and sterilized with a 20% (v/v) commercial bleach solution (Kao Ltd., Japan) for 30 minutes with gentle agitation. The rice seeds were then washed thoroughly with distilled water and placed on filter papers in 9cm petri dishes filled with 15 mL sterilized water in a 16-h light/8-h dark cycle at 26 °C for 1 week.

### *In vitro* infection assays with *Phtheirospermum*

Four to five days old *Phtheirospermum* seedlings were transferred for three days to nutrient-free 0.8% (w/v) agar medium or 0.8% (w/v) agar medium supplemented by nutrient or hormone treatment: ½MS, ½MS no N, 20.6 mM KNO_3_, 50 μM-20.6 mM NH_4_NO_3_, 0.62 mM KH_2_PO_4_, 1.9 mM NaH_2_PO_4_, 5 μM ABA, 10 μM Fluridone or 5 μM SA. Five day old *Arabidopsis* seedling were aligned next to and roots place in contact with these pre-treated *Phtheirospermum* roots for infection assays. Haustorium formation and xylem bridge development were measured at seven days post infection using a Zeiss Axioscope A1 microscope.

### Haustorium induction assay

Four to five days old *Phtheirospermum* seedlings were transferred to nutrient-free 0.8% (w/v) agar medium or 0.8% (w/v) agar medium supplemented by nutrients (½MS, ½MS no N, NH_4_NO_3_) for a three day pre-treatment. Subsequently, seedlings were transferred to 0.8% (w/v) agar medium containing DMBQ (Sigma-Aldrich) or DMBQ with or without nutrient treatment and grown vertically for four to five days for haustorium induction.

### Greenhouse experiments

Ten days old *Phtheirospermum* seedlings were germinated *in vitro* as described above. The seedlings were then transferred in pots with 50:50 soil:sand ratio. *Arabidopsis* seeds were sprinkled around the *Phtheirospermum* seedling. The pots were placed at 25°C and long day conditions (16-h light:8-h dark and 100 μmol m^-2^ s^-1^) and 60% humidity for 1.5 months. During this time the plants were given deionized water or water supplemented with fertilizer (commercial fertilizer Blömstra 51-10-43 N-P-K at 2 ml/L).

### Histological staining

Dissected roots were fixed in ethanol-acetic acid, stained with Safranin-O solution (0.1%), cleared with chloral hydrate for two to three days before observation with a Zeiss Axioscope A1 microscope as previously described (Cui et al., 2016).

### Xylem strand measurement

Five days old *Phtheirospermum* seedlings (n=19) were treated with 1 μM ABA or 5 μM ABA, ½MS no N, ½MS or 5 mM NH_4_NO_3_ for three days. Afterwards, the number of xylem strands were measured at 2 mm from the root tip with a Zeiss Axioscope A1 microscope.

### Sample preparation for RNAseq

40 four to five days old *Phtheirospermum* seedlings were transferred to nutrient-free 0.8% (w/v) agar medium or 0.8% (w/v) agar medium supplemented with 10.3 mM NH_4_NO_3_ or 0.08 μM BA for 3 days prior to infection with *Arabidopsis* Col-0. As a control group, 40 *Phtheirospermum* seedlings per treatment remained without the *Arabidopsis* host. For the water treatment samples, five time points were prepared (0,12,24,48,72 hpi). For the NH_4_NO_3_ and BA treatments, samples were prepared for three time points (0,12,24 hpi). One to two mm from *Phtheirospermum* and *Arabidopsis* root tips were harvested for the not infecting plants and the 0 hpi infecting plants. For the 12, 24, 48, 72 hpi time points, the haustorium, including 1-2 mm above and below tissue was collected together with the corresponding region of the *Arabidopsis* root. This experiment was replicated three times. RNA extraction was performed using the ROTI^®^Prep RNA MINI (Roth) kit following the manufacturer’s instructions. The isolation of mRNA and library preparation were performed using NEBNext^®^ Poly(A) mRNA Magnetic Isolation Module (#E7490), NEBNext^®^ Ultra™ RNA Library Prep Kit for Illumina^®^ (# E7530L), NEBNext^®^ Multiplex Oligos for Illumina^®^ (#E7600) following the manufacturer’s instructions. The libraries were then sequenced using paired end sequencing with an Illumina NovaSeq 6000.

### Bioinformatic analysis

The adapter and low-quality sequences were removed using the fastp software with default parameters (Chen et al., 2018). The quality-filtered reads were mapped to both the *Phtheirospermum* (Cui et al., 2020) and *Arabidopsis* genome (TAIR10) using STAR (Dobin et al., 2013) and were separated based on mapping to *Phtheirospermum* and *Arabidopsis* reads. The separated reads were then re-mapped to their respective genomes. The read count was calculated using FeatureCounts (Liao et al., 2014). The differential expression analysis was performed using Deseq2 (Love et al., 2014) (TableS2-S8). The gene expression clustering was performed using the Mfuzz software(Futschik et al., 2009). Custom annotations of the *Phtheirospermum* predicted proteins (Cui et al., 2020) were estimated using InterProScan (Blum et al., 2020), these were used for Gene ontology analysis that was performed using the topGO software (Alexa et al., 2016). ABA responsive, SA and cytokinin related genes in *Phtheirospermum* (TableS10) were identified using the tBLASTp and tBLASTp algorithm of the *Arabidopsis* ABA responsive genes described by (Nemhauser et al., 2006) or *Arabidopsis* genes involved in SA and CK pathways against the *Phtheirospermum* genome (Cui et al., 2020).

### Statistics

Statistical analyses were performed using ANOVA followed by Tukey’s HSD post-hoc test. For haustoria per *Phtheirospermum* and xylem bridge formation percentage data, the statistical analyses were performed on the means of at least 3 biological replicates, where each biological replicate consisted of 20 plants. For single comparisons, student’s t-tests were used.

### qPCR

*Phtheirospermum* seedlings were grown for five days before transferring to nutrient-free 0.8% (w/v) agar medium or 0.8% (w/v) agar medium supplemented with ½MS, ½MS no N or 5 μM ABA for 5 days. Additionally, *Phtheirospermum* seedlings were placed on pots containing 100:0, 50:50, 33:66, 25:75 soil:sand ratios. The pots were placed at 25°C and long day conditions (16 h light:8 h dark and 100 μmol m^-2^ s^-1^) and 60% humidity for 1.5 months. During this time the plants were provided deionized water. The seedlings or the shoots and roots of the above described *Phtheirospermum* were then harvested and RNA extraction was performed using the ROTI^®^Prep RNA MINI (Roth) kit following the manufacturer’s instructions. The extracted RNA was then treated with DNase I (Thermo Scientific™) following the manufacturer’s instructions. cDNA synthesis was performed using Maxima First Strand cDNA Synthesis Kit for RT-qPCR (Thermo Scientific™) following the manufacturer’s instructions. *PjPTB* (Ishida et al., 2016) was used as an internal control. qPCR was performed with SYBR-Green master mix (Applied Biosystems™). The relative expression was calculated using the Pfaffl method (Pfaffl, 2001). All experiments were repeated at least three times with at least two technical replications each. For statistical analysis, the student’s t-test was used. The primers used for this experiment are listed in TableS9.

### Hormonal quantifications

*Phtheirospermum* seedlings were grown for four to five days before transferring to nutrient-free 0.8% (w/v) agar medium or 0.8% (w/v) agar medium supplemented with 10.3mM NH_4_NO_3_ for three days. *Arabidopsis* Col-0 or *aba2-1* was placed next to the *Phtheirospermum* seedlings and left for 10 days. *Phtheirospermum* seedlings without a host were used as “not infecting” control. After 10 days with or without the presence of a host, four to five entire *Phtheirospermum* seedlings per sample and four to five entire *Arabidopsis* seedlings per sample were collected. The samples were crushed to powder using liquid N with mortar and pestle. Samples were extracted, purified and analyzed according a previously published method (Šimura et al., 2018). Briefly, approx. 20 mg of frozen material per sample was homogenized and extracted in 1 mL of ice-cold 50% aqueous acetonitrile (v/v) with the mixture of ^13^C- or deuterium-labelled internal standards using a bead mill (27 hz, 10 min, 4°C; MixerMill, Retsch GmbH, Haan, Germany) and sonicator (3 min, 4°C; Ultrasonic bath P 310 H, Elma, Germany). After centrifugation (14 000 RPM, 15 min, 4°C), the supernatant was purified as following. A solid-phase extraction column Oasis HLB (30 mg 1 cc, Waters Inc., Milford, MA, USA) was conditioned with 1ml of 100% methanol and 1ml of deionized water (Milli-Q, Merck Millipore, Burlington, MA, USA). After the conditioning steps each sample was loaded on SPE column and flow-through fraction was collected together with the elution fraction 1ml 30% aqueous acetonitrile (v/v). Samples were evaporated to dryness using speed vac (SpeedVac SPD111V, Thermo Scientific, Waltham, MA, USA). Prior LC-MS analysis, samples were dissolved in 40 μL of 30% acetonitrile (v/v) and transferred to insert-equipped vials. Mass spectrometry analysis of targeted compounds was performed by an UHPLC-ESI-MS/MS system comprising of a 1290 Infinity Binary LC System coupled to a 6490 Triple Quad LC/MS System with Jet Stream and Dual Ion Funnel technologies (Agilent Technologies, Santa Clara, CA, USA). The quantification was carried out in Agilent MassHunter Workstation Software Quantitative (Agilent Technologies, Santa Clara, CA, USA).

### *Striga-rice* Infection in the Rhizotron System

The rice infection was performed in a rhizotron system as described previously (Yoshida and Shirasu, 2009). Briefly, 7-d-old rice seedlings were transferred to the rhizotron (10-cm × 14-cm-square petri dish with top and bottom perforation for shoot growth and water draining, filled with same size of rockwool [Nichiasu, Tokyo, Japan] onto which a 100μm nylon mesh) and fertilized with 25 mL half-strength Murashige & Skoog media per rhizotron. The root parts of the rhizotron were covered with aluminum foil and placed vertically in a growth chamber at 12-h light: 28 °C /12-h dark: 20 °C cycles for two weeks before Striga infection. Rice seedlings were inoculated with *S. hermonthica* seeds by placing Strigol-treated *S. hermonthica* carefully along rice roots with 5 mm intervals. The rhizotron containing inoculated rice seedlings were incubated in the growth chamber described above, and developmental stages of *S. hermonthica* were categorized with a stereomicroscope (Zeiss Stemi 2000-C) after two and four weeks. Successful infection rates were calculated by the number of *Striga* with more than three leaf pairs divided by the total infected *Striga* seeds. Each rhizotron was watered with 25 mL of indicated nutrient or chemical containing solutions two times per week. The chemical concentrations used in this study was as followings; 10.3 mM ammonium nitrate, 1.09 mM monosodium phosphate, 20.6 mM potassium nitrate, 0.62 mM monopotassium phosphate, 10 μM gibberellic acid, 0.08 nM 6-benzylaminopurine, 10 μM paclobutrazol, 10 or 100 μM fluridone, and 10 or 100 μM abscisic acid.

### Accession numbers

Sequence data are available at the Gene Expression Omnibus (http://www.ncbi.nlm.nih.gov/geo/) under accession numbers GSE177484. Sequence data of the *Phtheirospermum japonicum* genes studied in this article are available in GenBank (http://www.ncbi.nlm.nih.gov/genbank/) under the accession numbers provided in TableS11.

## Supplemental tables

Supplemental figure S1: Nutrient availability does not affect Phtheirospermum shoot growth and xylem plate size

Supplemental figure S2: Gene ontology of the co-expression clusters

Supplemental figure S3: NH4NO3 affects gene expression and xylem genes

Supplemental figure S4: Gene ontology analysis of the up and down regulated genes in nitrate and BA not infecting treatments

Supplemental figure S5: Expression changes of ABA related genes

Supplemental figure S6: Host ABA levels do not affect Phtheirospermum infection

Supplemental figure S7: Effect of ABA on Striga

Supplemental table S1: gene lists of co-expression clusters

Supplemental table S2: water infecting vs water not infecting Deseq2 results

Supplemental table S3: nitrate infecting vs nitrate not infecting Deseq2 results

Supplemental table S4: BA infecting vs BA not infecting Deseq2 results

Supplemental table S5: nitrate infecting vs water infecting Deseq2 results

Supplemental table S6: nitrate not infecting vs water not infecting Deseq2 results

Supplemental table S7: BA infecting vs water infecting Deseq2 results

Supplemental table S8: normalized counts for Pj genes

Supplemental table S9: list of primers

Supplemental table S10: gene lists used in heatmaps

Supplemental table S11: accession numbers

## Author contributions

AK and CWM conceived the experiments. AK, ML, XZ and JS performed the experiments. SC, SY, KL and CWM supervised the experiments. AK and CWM wrote the paper. All authors edited and revised the final paper.

## Acknowledgements

We thank Annelie Carlsbecker and Prashanth Ramachandran for providing *abi1-1C* seeds, and Thomas Spallek for critical reading of the manuscript. AK, ML and CWM were supported by a Wallenberg Academy Fellowship (2016-0274) and an ERC starting grant (GRASP-805094). KL and JS were supported by grants from the Swedish Research Council, the Swedish Governmental Agency for Innovation Systems and the Knut and Alice Wallenberg Foundation. The authors acknowledge support from the Uppsala Multidisciplinary Center for Advanced Computational Science for assistance with access to the UPPMAX computational infrastructure. We also thank the Swedish Metabolomics Centre for access to instrumentation.

