## Supplemental Figures 1-7 for "Nitrates increase abscisic acid levels to regulate haustoria formation in the parasitic plant Phtheirospermum japonicum"

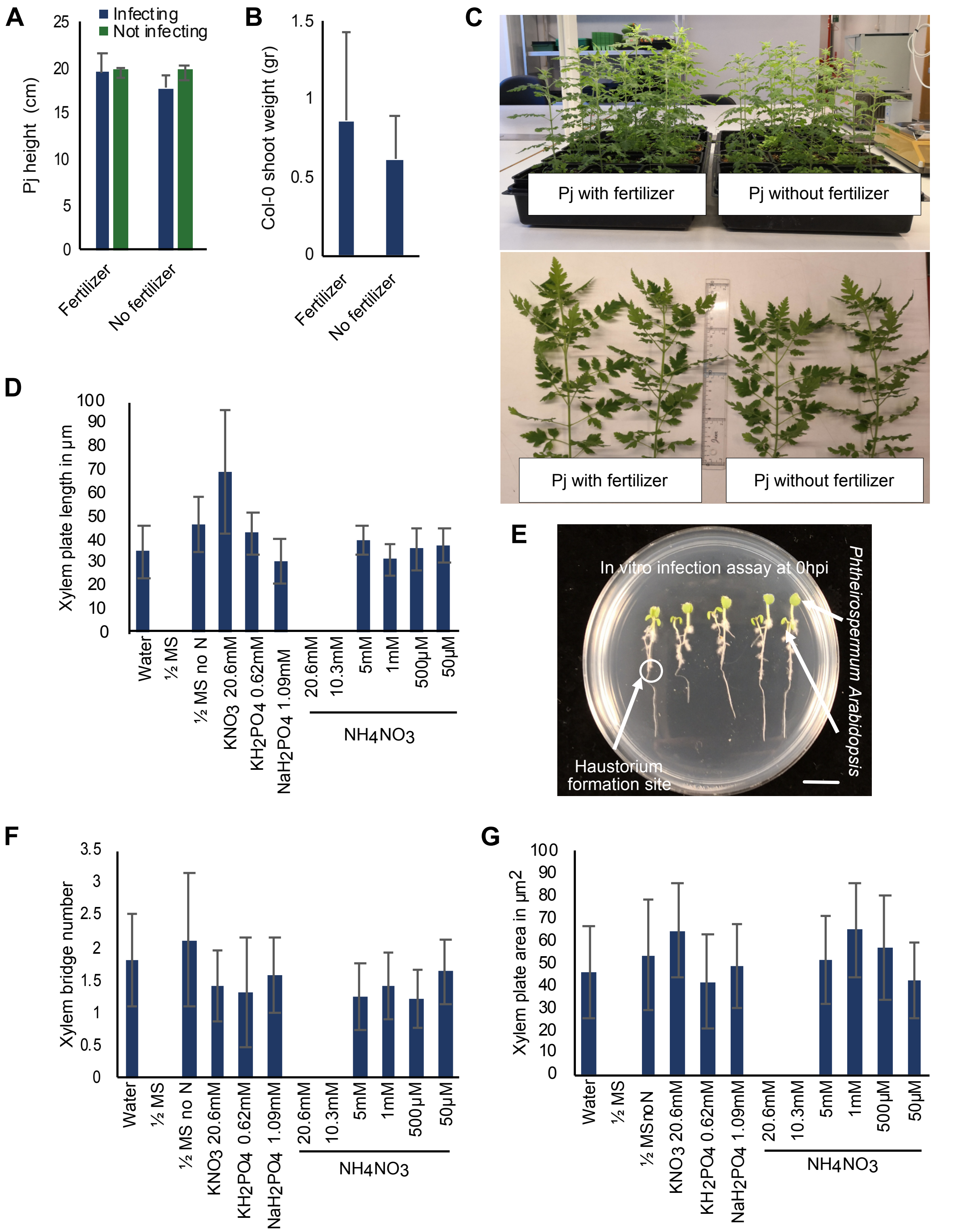


**Fig.S1 Nutrient availability does not affect *Phtheirospermum* shoot growth and xylem plate size.** (A-B) *Phtheirospermum* height and *Arabidopsis* Col-0 shoot weight with or without fertilizer application during infection of *Arabidopsis* by *Phtheirospermum*. (C) Photos of *Phtheirospermum* infecting with and without fertilizer application. (D-F-G) Xylem plate length (µM), xylem plate area (µm²) and xylem bridge number per haustorium under nutrient treatments (half strength MS, half strength MS no N, 20.6 mM $KNO_3$, 50 µM to 20.6 mM $NH_4NO_3$, 0.62 mM $KH_2PO_4$ or 1.9 mM $NaH_2PO_4$). (E) Photo of the *in vitro* infection assay set-up. (A,B,D,F,G) Bars represent mean ± SD. Scale bar 1 cm for (E).

**A**

| cluster 1 | | p-value |
|---|---|---|
| GO:0016192 | vesicle-mediated transport | 2.00E-06 |
| GO:0016567 | protein ubiquitination | 0.00023 |
| GO:0034613 | cellular protein localization | 0.00172 |
| GO:0023052 | signalling | 0.00263 |
| GO:0015693 | magnesium ion transport | 0.01378 |
| GO:0006808 | regulation of nitrogen utilization | 0.03685 |

| cluster2 | | p-value |
|---|---|---|
| GO:0006260 | DNA replication | 0.00022 |
| GO:0006397 | mRNA processing | 0.00045 |
| GO:0008654 | phospholipid biosynthetic process | 0.00079 |
| GO:0006281 | DNA repair | 0.00098 |
| GO:0006413 | translational initiation | 0.00594 |
| GO:0009611 | response to wounding | 0.00841 |
| GO:1901657 | glycosyl compound metabolic process | 0.01327 |
| GO:0046039 | GTP metabolic process | 0.01748 |
| GO:0006396 | RNA processing | 0.01842 |
| GO:0006366 | transcription by RNA polymerase II | 0.03476 |
| GO:0030243 | cellulose metabolic process | 0.03904 |

| cluster3 | | p-value |
|---|---|---|
| GO:0007018 | microtubule-based movement | 3.80E-13 |
| GO:0045944 | positive regulation of transcription | 2.90E-05 |
| GO:0000165 | MAPK cascade | 0.01294 |
| GO:0033014 | tetrapyrrole biosynthetic process | 0.03379 |
| GO:0007017 | microtubule-based process | 0.03922 |
| GO:0044267 | cellular protein metabolic process | 0.04835 |

| cluster4 | | p-value |
|---|---|---|
| GO:0006508 | proteolysis | 5.90E-08 |
| GO:2001141 | regulation of RNA biosynthetic process | 2.80E-05 |
| GO:0005975 | carbohydrate metabolic process | 0.00016 |
| GO:0005985 | sucrose metabolic process | 0.00488 |
| GO:0010411 | xyloglucan metabolic process | 0.00569 |
| GO:0046274 | lignin catabolic process | 0.00705 |
| GO:0007064 | mitotic sister chromatid cohesion | 0.01867 |
| GO:0006952 | defence response | 0.04476 |

| cluster 5 | | p-value |
|---|---|---|
| GO:0055114 | oxidation-reduction process | 6.00E-06 |
| GO:0009060 | aerobic respiration | 3.80E-05 |
| GO:0006979 | response to oxidative stress | 0.0024 |
| GO:0006561 | proline biosynthetic process | 0.0029 |
| GO:0042744 | hydrogen peroxide catabolic process | 0.0178 |
| GO:0009073 | aromatic amino acid family biosynthetic process | 0.0238 |
| GO:0030418 | nicotianamine biosynthetic process | 0.024 |
| GO:0045944 | positive regulation of transcription | 0.0281 |
| GO:0006633 | fatty acid biosynthetic process | 0.0293 |

| cluster6 | | p-value |
|---|---|---|
| GO:0016192 | vesicle-mediated transport | 3.30E-17 |
| GO:0046907 | intracellular transport | 3.80E-12 |
| GO:0034613 | cellular protein localization | 3.50E-11 |
| GO:0007264 | small GTPase mediated signal transduction | 1.00E-06 |
| GO:0030163 | protein catabolic process | 0.00015 |
| GO:0051225 | spindle assembly | 0.00017 |
| GO:0015991 | ATP hydrolysis coupled proton transport | 0.0005 |
| GO:0072330 | monocarboxylic acid biosynthetic process | 0.00088 |
| GO:0016310 | phosphorylation | 0.0013 |
| GO:0015833 | peptide transport | 2.40E-11 |
| GO:0016052 | carbohydrate catabolic process | 0.02928 |

| cluster 7 | | p-value |
|---|---|---|
| GO:0055114 | oxidation-reduction process | 0.00033 |
| GO:0006779 | porphyrin-containing compound biosynthesis | 0.00046 |
| GO:0030001 | metal ion transport | 0.00276 |
| GO:0006749 | glutathione metabolic process | 0.00666 |
| GO:0055085 | transmembrane transport | 0.00741 |
| GO:0051252 | regulation of RNA metabolic process | 0.02586 |
| GO:0009690 | cytokinin metabolic process | 0.02798 |
| GO:0006979 | response to oxidative stress | 0.03193 |

| cluster8 | | p-value |
|---|---|---|
| GO:0006412 | translation | < 1e-30 |
| GO:0022613 | ribonucleoprotein complex biogenesis | 2.10E-20 |
| GO:0006396 | RNA processing | 1.10E-13 |
| GO:0034660 | ncRNA metabolic process | 5.50E-09 |
| GO:0009451 | RNA modification | 5.40E-08 |
| GO:0006457 | protein folding | 1.10E-07 |
| GO:0032259 | methylation | 1.30E-07 |
| GO:0071826 | ribonucleoprotein complex subunit | 3.80E-07 |
| GO:0016071 | mRNA metabolic process | 7.10E-07 |
| GO:0042455 | ribonucleoside biosynthetic process | 1.50E-06 |
| GO:0009089 | lysine biosynthetic process | 3.60E-05 |

**Fig.S2 Gene ontology of the co-expression clusters.** (A) Gene ontology analysis for the differentially expressed genes assigned to each co-expression cluster. GO categories with the lowest p-values are shown ($P<0.05$).

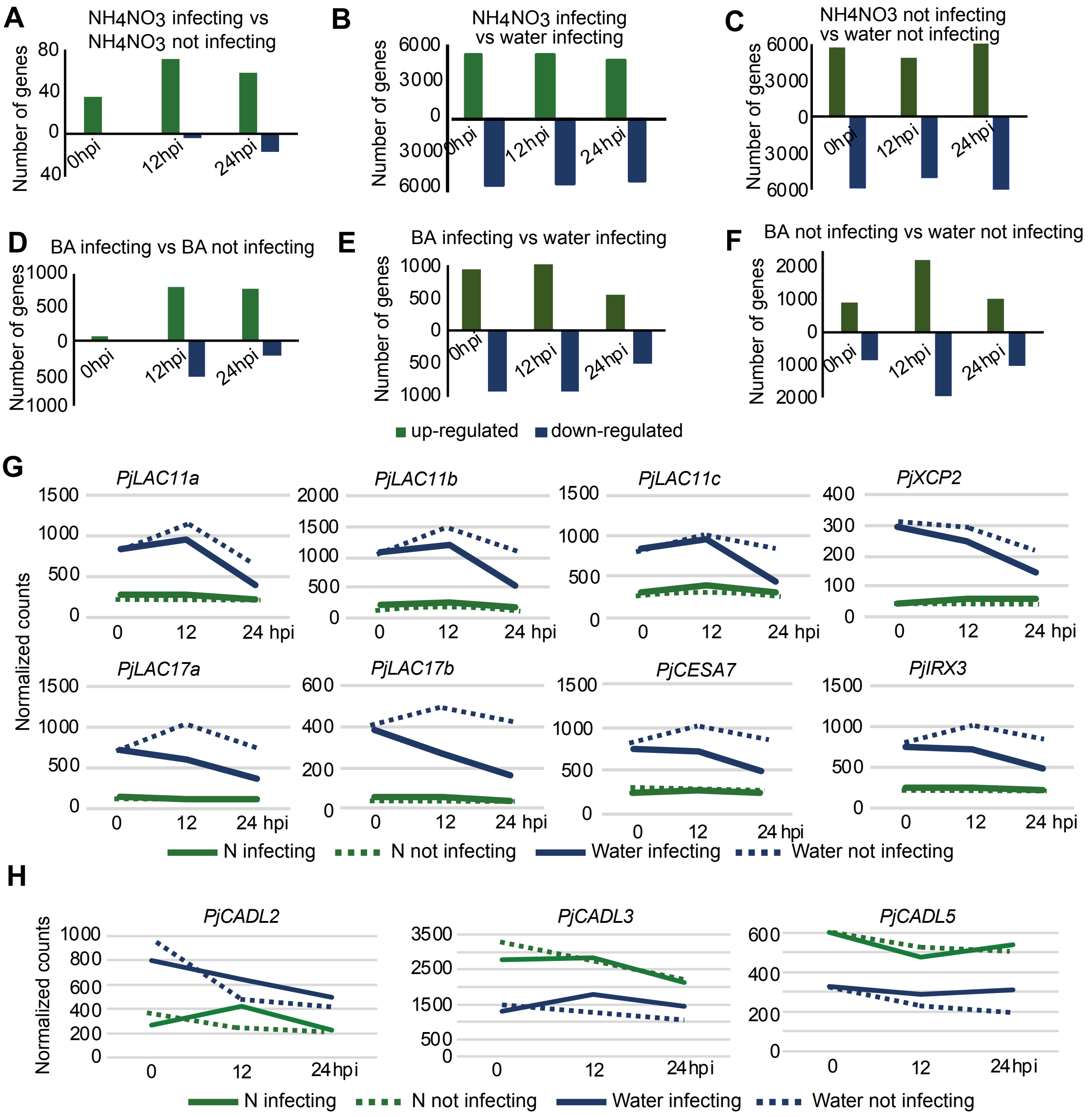


**Fig.S3 $NH_4NO_3$ affects gene expression and xylem genes.** (A-F) Number of genes differentially expressed over three time points in the $NH_4NO_3$ and BA RNAseq treatment between *Phtheirospermum* infecting and not infecting and in the $NH_4NO_3$ vs the water RNAseq treatment in *Phtheirospermum*. (G) Normalized counts of *PjXCP2, PjLAC11a,b,c, PjLAC17a,b, PjIRX3, PjCESA7* over three time points shown for *Phtheirospermum* infecting and not infecting in the $NH_4NO_3$ and water treatment. (H) Normalized counts of *PjCADL*2,3,5 over three time points shown for *Phtheirospermum* infecting and not infecting in the $NH_4NO_3$ and water treatment.

**A**

| nitrate not infecting vs water not infecting up-regulated | | | | | | | | |
|---|---|---|---|---|---|---|---|---|
| 0hpi | | p-value | 12hpi | | p-value | 24hpi | | p-value |
| GO:2000028 | regulation of photoperiodism | 0.0001 | GO:0008272 | sulfate transport | 0.0006 | GO:0009607 | response to biotic stimulus | 4.30E-07 |
| GO:0051252 | RNA metabolism regulation | 0.0011 | GO:0055114 | oxidation-reduction process | 0.0010 | GO:0051252 | RNA metabolism regulation | 1.70E-06 |
| GO:0006000 | fructose metabolic process | 0.0012 | GO:0051252 | RNA metabolism regulation | 0.0011 | GO:0055114 | oxidation-reduction process | 8.30E-05 |
| GO:0005992 | trehalose biosynthetic process | 0.0021 | GO:0005992 | trehalose biosynthesis | 0.0013 | GO:0019344 | cysteine biosynthetic process | 0.0003 |
| GO:0015698 | inorganic anion transport | 0.0022 | GO:0009584 | detection of visible light | 0.0017 | GO:0006952 | defense response | 0.0011 |
| GO:0015969 | guanosine tetraphosphate metabolism | 0.0023 | GO:0018298 | protein-chromophore linkage | 0.0017 | GO:0006563 | L-serine metabolic process | 0.0013 |
| GO:0055114 | oxidation-reduction process | 0.0032 | GO:0009607 | response to biotic stimulus | 0.0040 | GO:0019318 | hexose metabolic process | 0.0015 |
| GO:0015995 | chlorophyll biosynthesis | 0.0086 | GO:0006001 | fructose catabolic process | 0.0117 | GO:0009785 | blue light signaling pathway | 0.0038 |
| GO:0015979 | photosynthesis | 0.0087 | GO:0034755 | iron ion transmembrane transport | 0.0117 | GO:0006817 | phosphate ion transport | 0.0041 |
| GO:0000160 | phosphorelay signal transduction | 0.0100 | GO:0042128 | nitrate assimilation | 0.0117 | GO:0015969 | guanosine tetraphosphate metabolism | 0.0041 |

| nitrate not infecting vs water not infecting down-regulated | | | | | | | | |
|---|---|---|---|---|---|---|---|---|
| 0hpi | | p-value | 12hpi | | p-value | 24hpi | | p-value |
| GO:0042737 | drug catabolic process | 8.40E-06 | GO:0046274 | lignin catabolic process | 5.60E-08 | GO:0007017 | microtubule-based process | 4.30E-08 |
| GO:0046274 | lignin catabolic process | 9.50E-05 | GO:0042737 | drug catabolic process | 3.20E-06 | GO:0046274 | lignin catabolic process | 9.10E-05 |
| GO:0071554 | cell wall organization/biogenesis | 1.40E-03 | GO:0055114 | oxidation-reduction process | 7.30E-04 | GO:0008610 | lipid biosynthetic process | 3.20E-04 |
| GO:0006857 | oligopeptide transport | 0.0031 | GO:0006979 | response to oxidative stress | 0.0008 | GO:0009690 | cytokinin metabolic process | 0.0011 |
| GO:0006979 | response to oxidative stress | 0.0035 | GO:0005975 | carbohydrate metabolism | 0.0021 | GO:0006631 | fatty acid metabolic process | 0.0011 |
| GO:0055085 | transmembrane transport | 0.0124 | GO:0045492 | xylan biosynthetic process | 0.0057 | GO:0009813 | flavonoid biosynthesis | 0.0026 |
| GO:0005975 | carbohydrate metabolism | 0.0169 | GO:0007017 | microtubule-based process | 0.0066 | GO:0042737 | drug catabolic process | 0.0035 |
| GO:0009269 | response to desiccation | 0.0269 | GO:0000079 | regulation of protein synthesis | 0.0082 | GO:0008202 | steroid metabolic process | 0.0061 |
| GO:0044264 | polysaccharide metabolism | 0.0322 | GO:0015743 | malate transport | 0.0086 | GO:0055114 | oxidation-reduction process | 0.0062 |
| GO:0042219 | amino acid catabolism | 0.0366 | GO:0006542 | glutamine biosynthesis | 0.0139 | GO:0006555 | methionine metabolism | 0.0084 |

| BA not infecting vs water not infecting up-regulated | | | | | | | | |
|---|---|---|---|---|---|---|---|---|
| 0hpi | | p-value | 12hpi | | p-value | 24hpi | | p-value |
| GO:0009690 | cytokinin metabolic process | 1.50E-08 | GO:0006260 | DNA replication | 1.10E-26 | GO:0006412 | translation | 6.40E-07 |
| GO:0000160 | phosphorelay signal transduction | 8.60E-08 | GO:0006412 | translation | 2.60E-26 | GO:0009690 | cytokinin metabolic process | 6.60E-05 |
| GO:0055114 | oxidation-reduction process | 5.10E-06 | GO:0006457 | protein folding | 6.00E-16 | GO:0006457 | protein folding | 1.30E-04 |
| GO:0006006 | glucose metabolic process | 0.0001 | GO:0007018 | microtubule-based movement | 0.0000 | GO:0009152 | purine ribonucleotide synthesis | 0.0004 |
| GO:0006855 | drug transmembrane transport | 0.0004 | GO:0006265 | DNA topological change | 0.0001 | GO:0009168 | ribonucleoside monophosphate synth[illegible] | 0.0004 |
| GO:0009607 | response to biotic stimulus | 0.0027 | GO:0006310 | DNA recombination | 0.0001 | GO:0009206 | ribonucleoside triphosphate synthesis | 0.0005 |
| GO:0042737 | drug catabolic process | 0.0057 | GO:0009690 | cytokinin metabolic process | 0.0001 | GO:0006855 | drug transmembrane transp. | 0.0008 |
| GO:0008272 | sulfate transport | 0.0101 | GO:0009082 | amino acid biosynthesis | 0.0003 | GO:0006979 | response to oxidative stress | 0.0025 |
| GO:0009664 | plant-type cell wall organization | 0.0177 | GO:0006281 | DNA repair | 0.0008 | GO:0042744 | hydrogen peroxide catabolism | 0.0027 |
| GO:0006979 | response to oxidative stress | 0.0200 | GO:0065004 | protein-DNA complex assembly | 0.0014 | GO:0009664 | cell wall organization | 0.0048 |

| BA not infecting vs water not infecting up-regulated | | | | | | | | |
|---|---|---|---|---|---|---|---|---|
| down-regulated | | p-value | down-regulated | | p-value | down-regulated | | p-value |
| GO:0019419 | sulfate reduction | 0.00076 | GO:0055114 | oxidation-reduction process | 7.10E-05 | GO:0006355 | regulation of transcription | 4.60E-04 |
| GO:0071555 | cell wall organization | 0.00275 | GO:0046274 | lignin catabolic process | 7.70E-05 | GO:0006857 | oligopeptide transport | 5.90E-04 |
| GO:0006857 | oligopeptide transport | 0.00332 | GO:0019344 | cysteine biosynthesis | 3.70E-04 | GO:0019419 | sulfate reduction | 9.60E-04 |
| GO:0048544 | recognition of pollen | 0.00918 | GO:0006468 | protein phosphorylation | 0.0005 | GO:0005992 | trehalose biosynthesis | 0.0024 |
| GO:0046274 | lignin catabolic process | 0.01246 | GO:0009607 | response to biotic stimulus | 0.0006 | GO:0019344 | cysteine biosynthesis | 0.0041 |
| GO:0055085 | transmembrane transport | 0.01424 | GO:0005992 | trehalose biosynthesis | 0.0009 | GO:0055085 | transmembrane transport | 0.0072 |
| GO:0015936 | coenzyme A metabolism | 0.01453 | GO:0051179 | localization | 0.0011 | GO:0048544 | recognition of pollen | 0.0147 |
| GO:0006270 | DNA replication initiation | 0.02401 | GO:0042737 | drug catabolic process | 0.0013 | GO:0055114 | oxidation-reduction process | 0.0223 |
| GO:0070588 | Ca ion transmembrane transp. | 0.02401 | GO:0070588 | Ca ion transmembrane transp. | 0.0017 | GO:0016567 | protein ubiquitination | 0.0286 |
| GO:0055114 | oxidation-reduction process | 0.02517 | GO:0008283 | cell proliferation | 0.0024 | GO:0006558 | L-phenylalanine metabolism | 0.0299 |

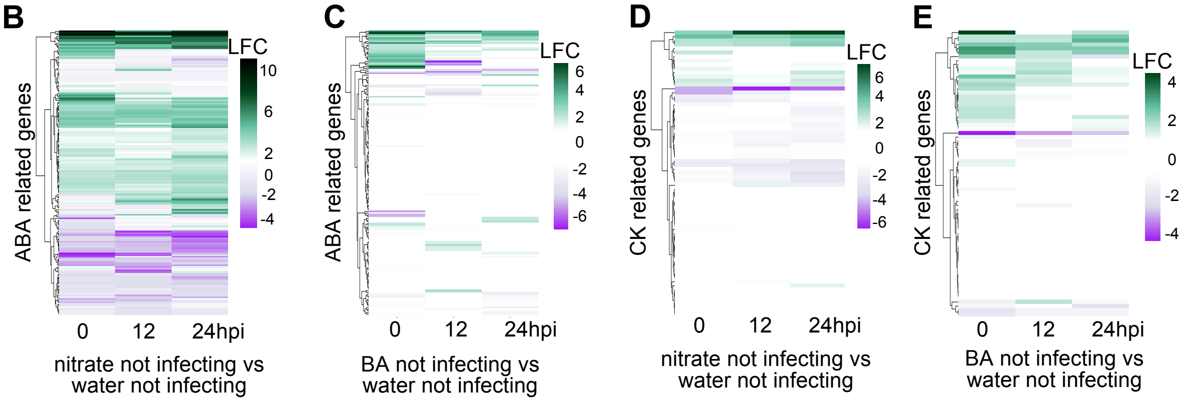


**Fig.S4 Gene ontology analysis of the up and down regulated genes in nitrate and BA not infecting treatments.** (A) Gene ontology analysis for genes differentially expressed during the three time points in the $NH_4NO_3$ not infecting and BA not infecting vs water not infecting RNAseq treatments in *Phtheirospermum*, shown are the top 10 GO categories with P< 0.05. (B-E) Heatmaps of the log2 fold change of 170 genes homologous to *Arabidopsis* ABA responsive genes and 72 cytokinin related genes shown over three time points in the water infecting vs water not infecting or in the $NH_4NO_3$ not infecting vs water not infecting and BA not infecting vs water not infecting RNAseq treatments in *Phtheirospermum*.

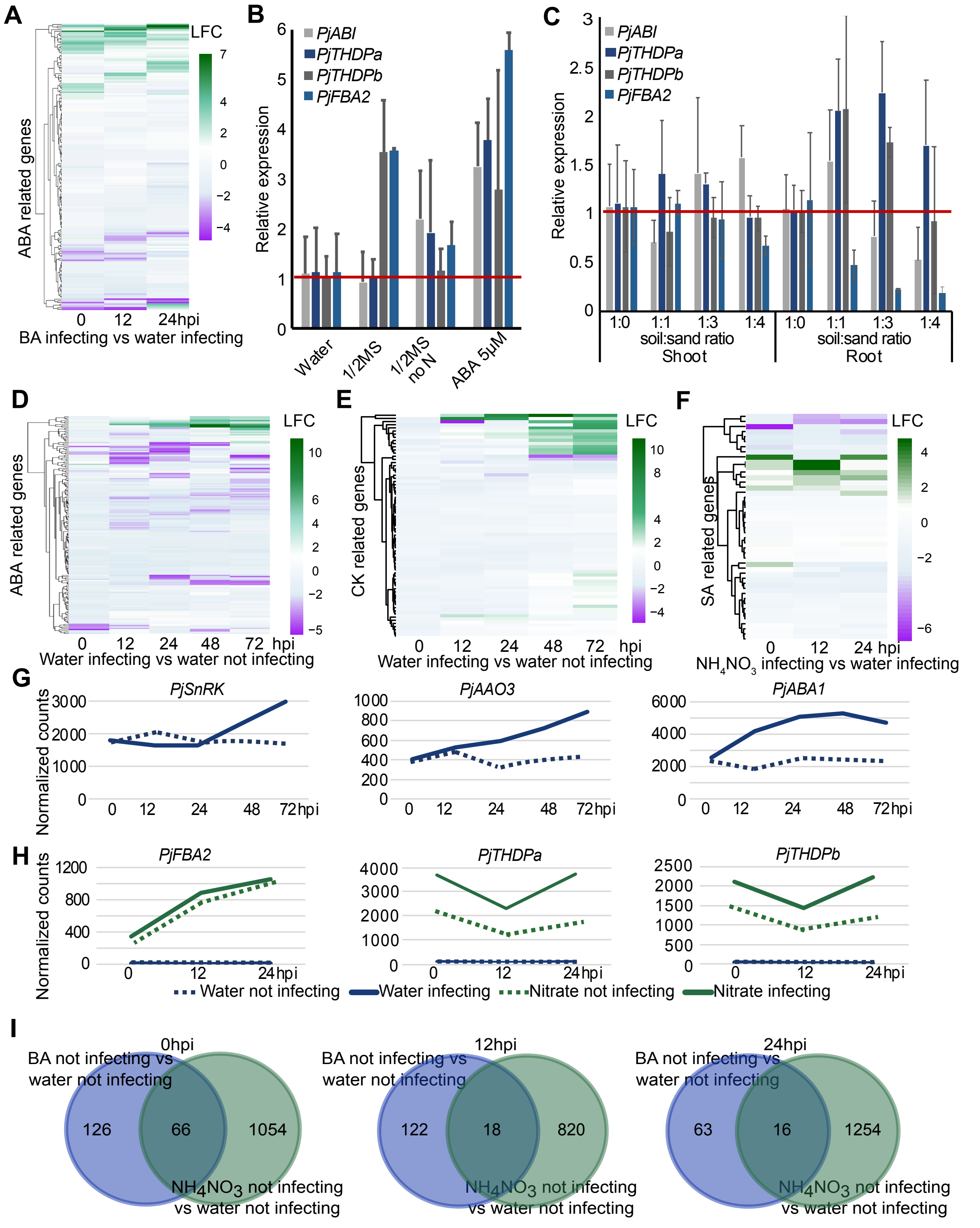


**Fig.S5 Expression changes of ABA related genes.** (A) Heatmap of the log2 fold change of 170 genes homologous to *Arabidopsis* ABA responsive genes shown over three time points in the BA infecting vs water infecting RNAseq treatment in *Phtheirospermum*. (B-C) Expression levels of *PjABI, PjFBA2 and PjTHDPa,b* under ½ MS, ½ MS no N and 5 μM ABA and various soil:sand ratios analyzed by RT-qPCR. (D-E-F) Heatmaps of the log2 fold change of 170 genes homologous to *Arabidopsis* ABA responsive genes, 72 cytokinin related genes and 45 SA related genes shown over five time points in the water infecting vs water not infecting or over three time points in the $NH_4NO_3$ infecting vs water infecting RNAseq treatments in *Phtheirospermum.* (G-H) Normalized counts of *PjSnRK, PjAAO3, PjABA1, PjFBA2, PjTHDPa,b* over five time points shown for *Phtheirospermum* water infecting and water not infecting and over three time points in the $NH_4NO_3$ infecting or $NH_4NO_3$ not infecting. (I) Venn diagrams of the differentially expressed genes in BA and nitrate not infecting over three time points in *Phtheirospermum*.

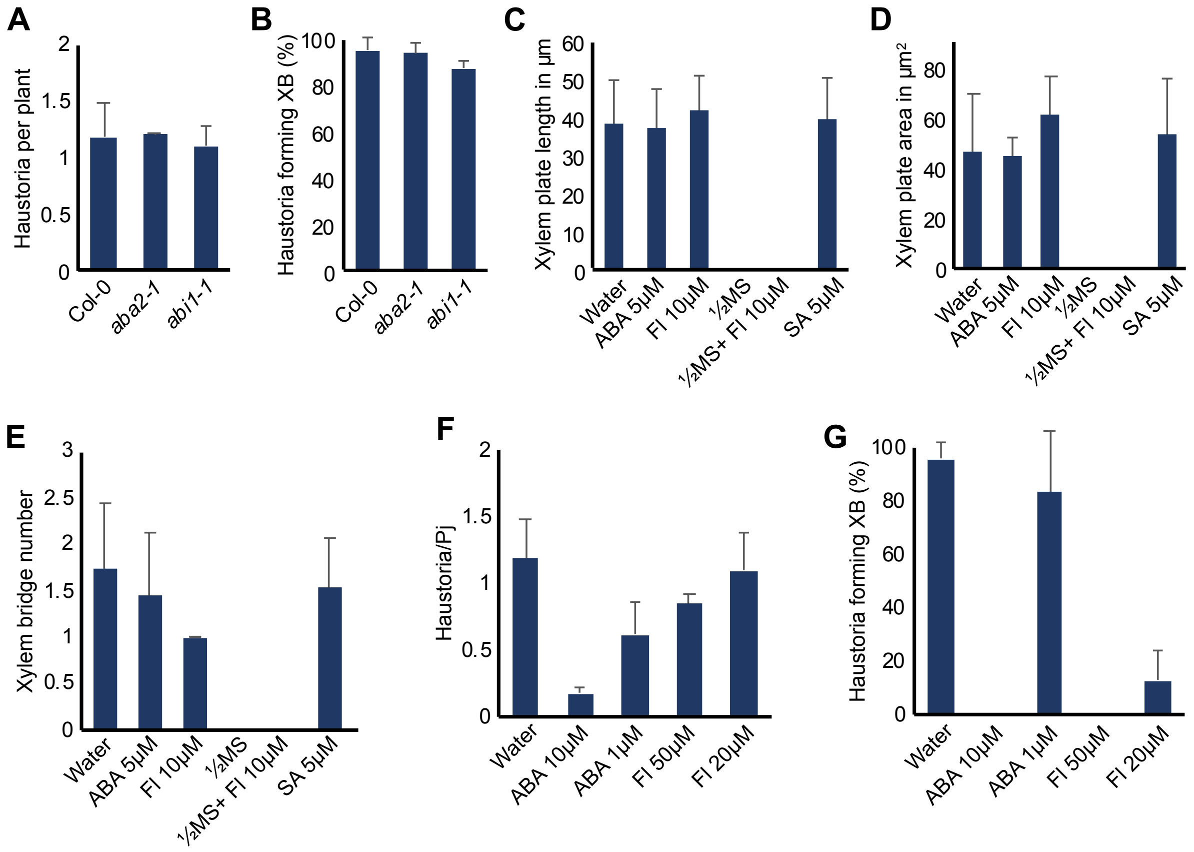


**Fig.S6 Host ABA levels do not affect *Phtheirospermum* infection.** (A-B) Average number of haustoria per *Phtheirospermum* seedling and xylem bridge formation percentage in *in vitro* infection assays with *Arabidopsis* Col-0, *aba2-1* and *abi1-1C* (*abi1-1*) as the host. (C-D-E) Xylem plate length (µM), xylem plate area (µm$^2$) and xylem bridge number per haustorium under 5 µM ABA, 10 µM fluridone, ½MS, ½MS + 10 µM fluridone or 5 µM SA treatments. (F- G) Average number of haustoria per *Phtheirospermum* seedling and xylem bridge formation percentage in *in vitro* infection assays with *Arabidopsis* Col-0 under 5 µM ABA, 10 µM fluridone, 5 mM $NH_4NO_3$, 5 mM $NH_4NO_3$ + 10 µM fluridone or 5 µM SA treatment. Bars represent mean ± SD.

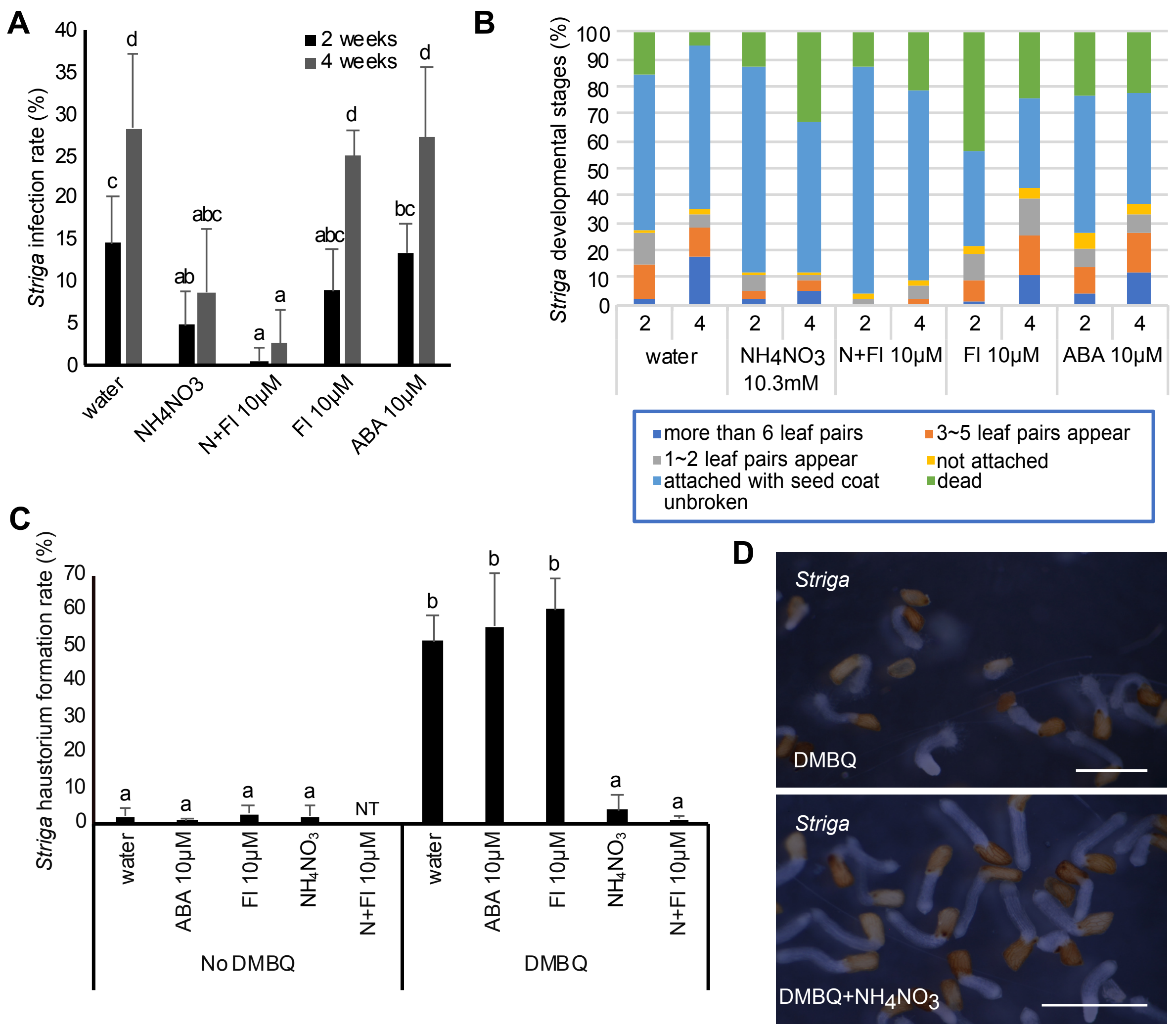


**Fig.S7 Effect of ABA on *Striga*.** (A) *Striga* infection rates at two and four weeks after infection with rice as a host under 5 μM ABA, 10 μM fluridone, 5 mM $NH_4NO_3$ or 5 mM $NH_4NO_3$ + 10 μM fluridone treatments. (B) Effect of 5 μM ABA, 10 μM fluridone, 5 mM $NH_4NO_3$ or 5 mM $NH_4NO_3$ + 10 μM fluridone treatments on *Striga* development at two and four weeks after infection. (C) Effect of 5 μM ABA, 10 μM fluridone, 5 mM $NH_4NO_3$ or 5 mM $NH_4NO_3$ + 10 μM fluridone treatment on *Striga* haustorium induction by 1 μM DMBQ. (D) Brightfield images of *Striga* haustorium formation assay with 1 μM DMBQ or 1 μM DMBQ+ 10.3 mM $NH_4NO_3$ at 1 day after treatment. (A, C) Bars represent mean ± SD (ANOVA $P<0.05$). Scale bars 1 mm for (D).
